# Interactive design of GPU-accelerated Image Data Flow Graphs and cross-platform deployment using multi-lingual code generation

**DOI:** 10.1101/2020.11.19.386565

**Authors:** Robert Haase, Akanksha Jain, Stéphane Rigaud, Daniela Vorkel, Pradeep Rajasekhar, Theresa Suckert, Talley J. Lambert, Juan Nunez-Iglesias, Daniel P. Poole, Pavel Tomancak, Eugene W. Myers

## Abstract

Modern life science relies heavily on fluorescent microscopy and subsequent quantitative bio-image analysis. The current rise of graphics processing units (GPUs) in the context of image processing enables batch processing large amounts of image data at unprecedented speed. In order to facilitate adoption of this technology in daily practice, we present an expert system based on the GPU-accelerated image processing library CLIJ: The CLIJ-assistant keeps track of which operations formed an image and suggests subsequent operations. It enables new ways of interaction with image data and image processing operations because its underlying GPU-accelerated image data flow graphs (IDFGs) allow changes to parameters of early processing steps and instantaneous visualization of their final results. Operations, their parameters and connections in the IDFG are stored at any point in time enabling the CLIJ-assistant to offer an undo-function for virtually unlimited rewinding parameter changes. Furthermore, to improve reproducibility of image data analysis workflows and interoperability with established image analysis platforms, the CLIJ-assistant can generate code from IDFGs in programming languages such as ImageJ Macro, Java, Jython, JavaScipt, Groovy, Python and C++ for later use in ImageJ, Fiji, Icy, Matlab, QuPath, Jupyter Notebooks and Napari. We demonstrate the CLIJ-assistant for processing image data in multiple scenarios to highlight its general applicability. The CLIJ-assistant is open source and available online: https://clij.github.io/assistant/

## 1 Introduction

The availability of image analysis algorithms exploiting the computational power of graphics processing units (GPUs) to a broader audience boosts the need for accessible tools for building GPU-accelerated image analysis workflows; in the life sciences and in adjacent imaging-dependent research fields. Typically, designing data analysis procedures utilizing GPUs involves expertise in programming and knowledge of GPU-specific programming languages such as the Open Computing Language (OpenCL) [1]. We demonstrate how one can construct complete image analysis workflows without writing a single line of OpenCL by assembling workflows from operations provided by the CLIJ framework [2]. We called the user interface CLIJ-asssistant because it allows interactive design of image data flow graphs (IDFGs) in ImageJ [3] or Fiji [4], while guiding the user with automatic suggestions. Depending on previously executed operations, it only shows operations that are suitable to the currently selected image, as shown in Figure 1. Automatic suggestions, semi-automated parameter optimization, and the immediate view of results enabled by GPU-acceleration allow rapid assembly of complex image analysis procedures on screen. IDFGs can also be stored to disc, reloaded, and version controlled. Furthermore, code-generation in multiple programming languages enables deployment of the same workflow to multiple collaborators working with different programming languages on different platforms.

**Figure 1:**
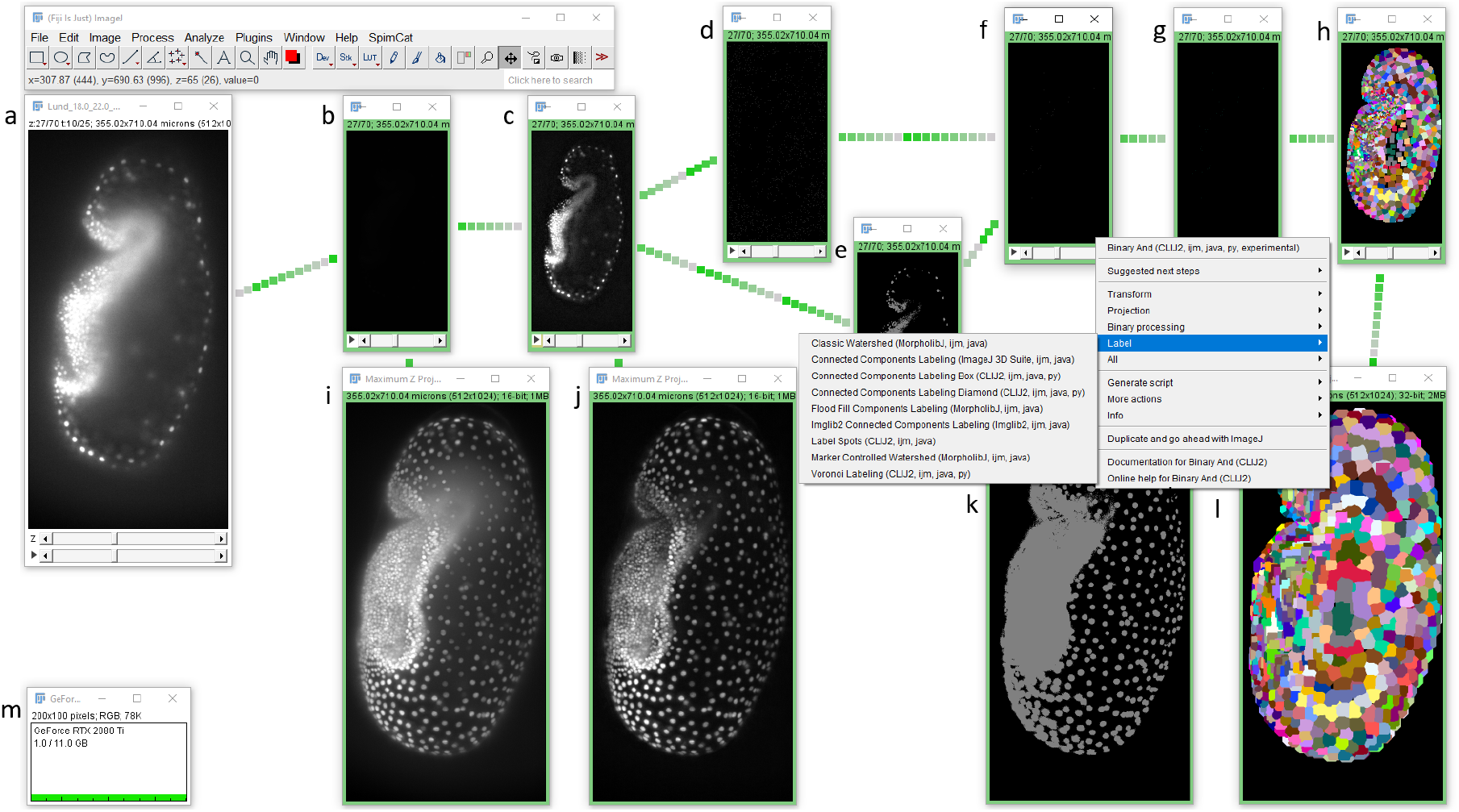
An IDFG allows assembly of image processing workflows and display of intermediate results in real-time. The presented graph takes a 4D image stack (a), pushes the current frame to the GPU memory (b), applies a top-hat filter for background subtraction (c), spot detection (d), thresholding (e), binary AND (f) spot labeling (g) and labeled spot extension (h). To facilitate examination of intermediate results, maximum intensity projections for intermediate results of pushing (i), background subtraction (j), thresholding (k) and label extension (l) are shown. The GPU memory display (m) allows to monitor available memory while setting up an IDFG. From any step in the graph, the user can select suitable subsequent operations using the shown right-click menu. The example shows the menu for the binary AND operation.

## 2 Methods

### Image data flow graph

We implemented the CLIJ-assistant under the hood of the established ImageJ user-interface so that ImageJ users can adopt it without changing their habits to much. In the background, CLIJ-assistant manages an image data flow graph (IDFG): Data flow graphs are a technical abstraction of processing workflows where graph nodes represent operations and connections between nodes, so called edges, represent data, which is sent from operation to successor operations. In our IDFGs, images are the edges as image data flows from operation to operation. Image windows on screen represent image processing operations and thus, the nodes of the graph. Technically, we implemented a directed acyclic graph. This means that image data are propagated in the graph in one direction only and no loops can be constructed. Basic usage of IDFGs is demonstrated in Supplementary Figure S1. Our approach is similar to the user-interfaces of Icy [5] and Knime [6]. An advantage is that intermediate results of the whole graph are always images which can be updated instantly in ImageJ’s user-interface while the user changes parameters. On the other hand, this limits available functionality compared to Knime and Icy data flow graphs: CLIJ-assistant focuses on image processing and analysis. In order to further optimize performance, the image stacks are kept in the GPU’s memory while minimizing pushing and pulling image data to/from GPU memory. Configuration dialogs of multiple steps can be open on screen. If the user changes a parameter, the affected operation and all subsequent operations are computed. In order to visualize the state of the graph, IDFG windows appear differently than standard ImageJ windows by their frame color, visualizing the graph’s execution state: A window with a red frame indicates that the shown image is invalid and will be computed as soon as intermediate results higher in the graph hierarchy are available. A yellow frame shows that computation is currently ongoing.

A green frame indicates that computation is finished. Furthermore, when moving a window, all downstream windows also move. This gives the user an impression of connections between graph nodes. This intuitive way of interacting with image data allows users to learn image processing, analysis and relationships between operations efficiently because it keeps technical implementation details out of sight.

For every processed image, the CLIJ-assistant can backtrack which original images, subsequent operations and corresponding parameters were involved in forming the image. Thus, it is not necessary to store intermediate results of various parameter configurations in the computer memory. The original image, the graph structure and the parameters of operations are sufficient to recompute a result. By storing the settings before every change, a history of settings is collected. The possibility of rewinding to former parameter settings brings a virtually unlimited undo functionality to Fiji. Users can select earlier parameter settings from a menu and get results back from former graph configurations, as shown in Supplementary Figure S2. Furthermore, users can save IDFGs to disc and reopen them in Fiji in a text file format. This allows application of version control systems to document parameter changes within and between specific projects.

### Available operations and extensibilty

After installation of the CLIJ-assistant, about 249 image processing operations are available. They include classic image filtering, spatial projections and transformations, namely rigid and affine transformations and image warping. For image segmentation, processing steps like thresholding, spot detection, binary post-processing and labeling operations can be chosen by the IDFG designer. Furthermore, machine learning tools for pixel classification and labelled object classification based on the Waikato Environment for Knowledge Analysis (Weka) [7] and the Trainable Weka Segmentation [8] can be assembled in the graph. An example workflow utilizing label classification is shown in Supplementary Figure S3. Tools for analyzing labelled objects, their neighbor relationships and topological parameters of adjacency graphs derived from images can be used as well.

The CLIJ-assistant uses the ImageJ2 plugin mechanism [9] to automatically discover additional CLIJ-compatible plugins. In that way, custom third-party operations can be introduced as graph nodes. These plugins do not necessarily need to be GPU-accelerated. To demonstrate the extensibility, about 50 additional image processing operations, utilizing the open-source libraries ImageJ, ImageJ2, Imglib2 [10], BoneJ [11], MorpholibJ [12], the ImageJ 3D Suite [13] and SimpleITK [14], are available for installation and testing via a separate Fiji update site.

### Expert system

State-of-the-art image analysis teaching courses introduce beginners to the concept of image analysis workflows as an assembly of operations. Concatenated operations stream input images via intermediate filtered images and regions of interest to quantitative measurements in arrays and tables. Typically, early processing steps are denoising, background removal, edge enhancement and image normalization. Afterwards, procedures in categories like segmentation, binarization, regionalization, detection and labeling follow. Finally, feature extraction and quantitative analysis follow to determine spatio-temporal properties of the imaged objects using descriptive statistics of pixel, region and topological properties. To facilitate accessibility and application of those steps, a search-bar and auto-completion of commands in the script editor were introduced in the recent years in Fiji. These tools allow the user to read in the graphical user-interface what certain operations do and how they can be applied to images. However, neither the search bar nor the auto-completion suggest what to do next. The user still needs to know the right terms to search for and thus, requires substantial knowledge of image analysis terminology. In order to overcome this limitation, we implemented an expert system in the CLIJ-assistant. Expert systems are an early form of artificial intelligence developed half a decade ago to model computationally for example the decision making processes clinicians used to reason under imperfect knowledge [15]. Our implementation of such an expert system guides users in choosing the right operations step-by-step by making context-dependend suggestions based on previously executed operation. The user interacts with it via the right-click menu of image windows as shown in Figure 1. For example, connected component labeling is typically applied after the image has been binarized, e.g. using Otsu’s thresholding method [16]. Since this connection is known, the CLIJ-assistant suggests connected component labeling after Otsu-thresholding. Proposed suggestions are derived from a knowledge base which consists of 1) typically subsequent operations and 2) technically compatible operations. The first part is derived from expert knowledge by extracting pairs of following operations from existing ImageJ macros. This knowledge base is delivered to end-users via the installation and update process. Furthermore, users can extend the local knowledge base by extracting additional pairs of subsequent operations from a local folder containing macro files. The knowledge base is a human-readable text file that can be exchanged and version control can be applied. With the underlying extraction process, experienced CLIJ-users can share their knowledge in an abstract representation without the need to disclose their potentially confidential image processing scripts. We encourage CLIJ power users to take part in the open call for contributions^1^ in order to combine knowledge from multiple experts and build up a community-driven knowledge base. The knowledge extraction and the deployment processes are both open-source and opt-in.

The second part of the knowledge base is implemented inside the CLIJ2 library and compatible libraries. It is based on the classification of operations and their inputs and output data types. For example, thresholding operations take images of any kind as input and produce binary images as an output. Thus, a category called ‘Binarize’ contains all operations with the same characteristic. Another ‘Labeling’ category holds operations that take images of any kind and assigns object-affiliation as pixel intensity values, also known as label images. The right-click menu only suggests operations that are applicable to the current image depending on the context as shown in Figure 2. Furthermore, when viewing the options for processing 2D images, no 3D-operations are shown and no 3D-to-2D projections. This reduces the number of suitable operations offered by the user-interface to a minimum, easing exploration of potential next processing steps.

**Figure 2:**
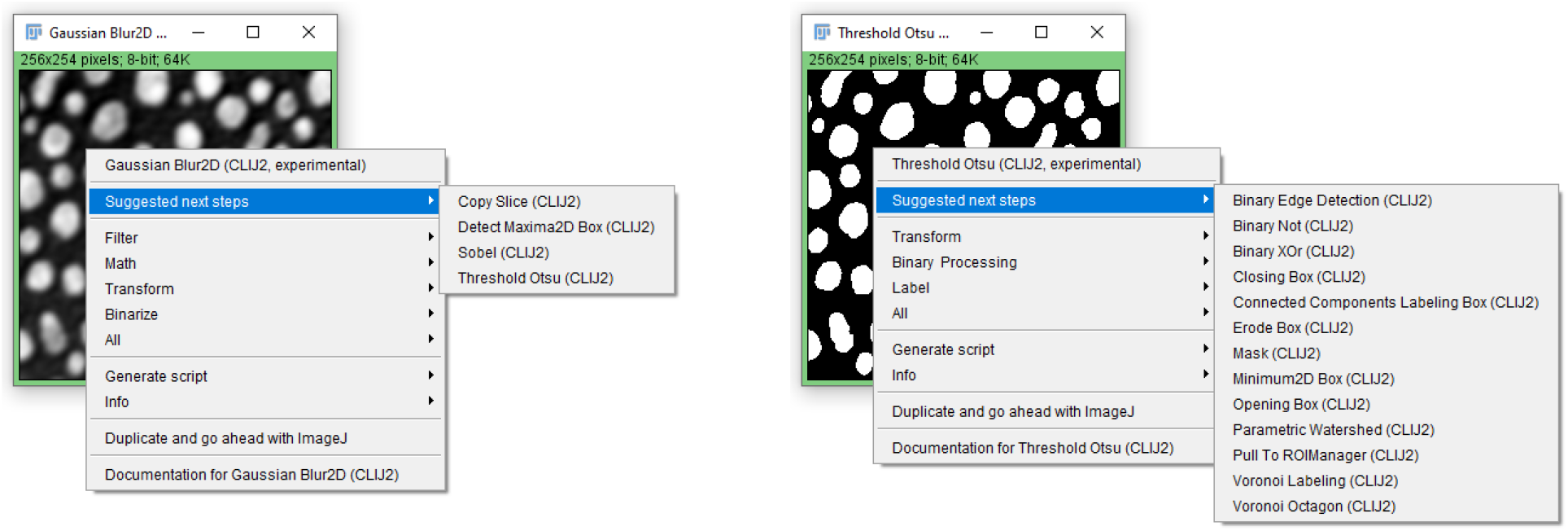
A built-in expert system suggests possible subsequent processing steps depending on the former steps applied in the workflow. Different operations are suggested when working with a grey value images (left) compared with a binary images (right).

If an expert applied a non-suitable operation in an earlier workflow, which was part of the knowledge-extraction process explained above, this operation will be suggested nevertheless. Furthermore, all operations are available under all circumstances from Fiji’s search bar giving the end-user full access to all available CLIJ-assistant compatible operations.

### Semi-automated parameter optimization

Professional image analysts set up data analysis workflows according to their experience. This can be straightforward if the expert has analyzed images of a similar kind before. Determining the optimal parameter configuration of such a workflow can be more complicated and is hindered by necessary additional efforts for determining the quality of results, for example if reference annotations are yet missing. Therefore, we introduce a simple annotation tool based on ImageJ’s ROI Manager and an automatic parameter optimization tool based on the simplex optimizer [17] and the gradient descent method implemented in the Apache Commons Math library [18]. For operations resulting in binary images, the parameter optimization can be used as demonstrated in Supplementary Figure S4. Therefore, all numeric parameters in the graph are considered for the optimization. The right-click menu also offers advanced options, for example to exclude parameters from the optimization process. As such a straightforward optimization approach may converge in a local optimum in parameter space, it is recommended to start the optimization with a good manual initial guess.

### Code generation for automation, documentation and knowledge exchange

The well-known Macro Recorder is one of the key features of ImageJ [3]. If the recorder is shown on screen while the user applies operations to images, it records code in ImageJ’s Macro programming language, which corresponds to the actions of the user. World-wide, a large number of users self-taught the principles of programming image analysis procedures by observing what is recorded in ImageJ. Also, the recording of object-oriented programming languages, such as JavaScript and Java, is available. Advanced programming skills are typically necessary to make these recorded scripts executable. Fiji [4] additionally introduced more advanced scripting languages such as Groovy, Jython, Clojure and Beanshell. However, macro recording capabilities for those scripting languages were not implemented. While the macro recorder contributed substantially to knowledge-transfer on how to use ImageJ Macro for automation of quantitative image analysis, adoption of the more modern and capable languages still has potential. To approach this aspect, we implemented code generation capabilities in the CLIJ-assistant using an IDFG as starting point. After an IDFG has been set up, configured and optimized, the graph can be exported in various programming languages. As the user chooses the language to export the IDFG to after setting it up, it is also possible to compare scripts in multiple languages as shown in Figure 3.

**Figure 3:**
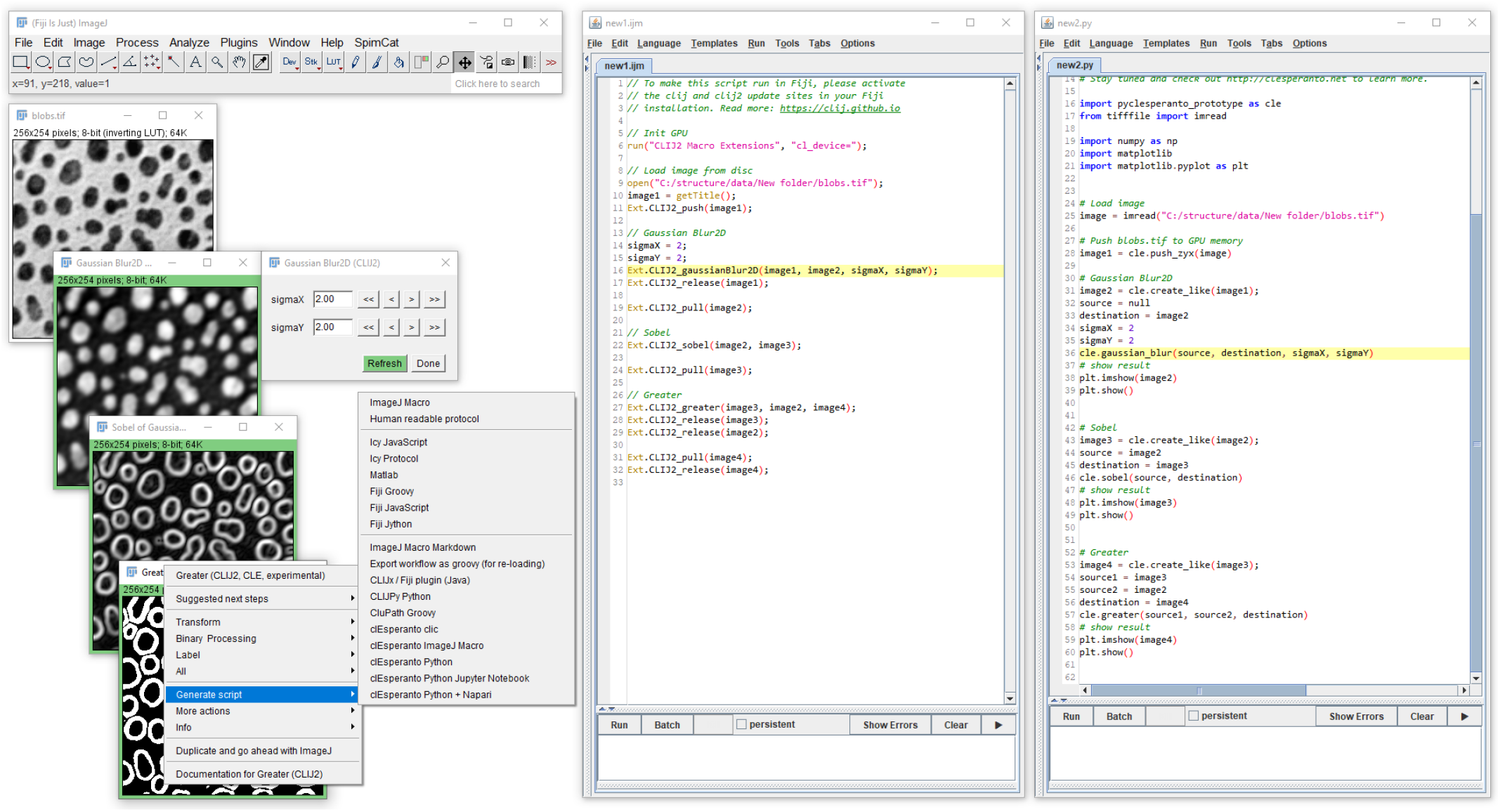
The IDFG shown on the left can be exported in multiple scripting languages. This enables users to compare scripts which execute the same workflow in different languages, for example in ImageJ Macro (center) and Python (right).

The exported scripts allow the user to go beyond ImageJ and Fiji, because they also offer programming languages applicable in other platforms. Currently supported languages are ImageJ Macro, Icy [5] Javascript, Matlab (Mathworks, United States MA), Fiji Groovy, ImageJ JavaScript, and Fiji Jython. Furthermore, a Java-plugin generator for Fiji, shown in Supplementary Figure S5, an Icy protocol generator, shown in Supplementary Figure S6, and script export for QuPath [19] Groovy, shown in Supplementary Figure S7, are available for testing. Support for Python [20] and C++ are under development and prototypes can be tested as well. Even though the set of implemented GPU-accelerated image processing functions available in Python and C++ are limited yet, we encourage early adopters to explore these technologies and provide feedback to guide further development. Through the introduced Python compatibility, the multi-dimensional image data explorer napari [21] and Jupyter notebooks [22] can be utilized as shown in Supplementary Figures S8 and S9, respectively. Interoperability is ensured with Python and C++ on a low code-level. In order to implement IDFG operations as similar as possible in ImageJ, in Python and in C++, the underlying GPU-accelerated OpenCL code is identical. Maintenance of the native code base is done as part of the clEsperanto^2^ project.

Fostering reproducibility of image analysis procedures by clear documentation, human-readable protocols of the IDFG can be exported as shown in Supplementary Figure S10. These protocols can be used to communicate the applied image processing workflow with scientists who are used to other platforms and to people without coding experience. Hence, we see this feature as a potential supplementary methods section generator. ImageJ Macro Markdown notebooks [23] can be generated from an IDFG as well. These notebooks present a hybrid between human-readable protocols and macro executable code, as shown in Supplementary Figure S11.

## 3 Results

The introduced toolkit for designing IDFGs allows to setup complex image processing routines without coding. To demonstrate the capabilities, we chose four typical use cases from gut neuroscience, developmental biology and cancer research. For all three example workflows, raw data and IDFGs can be found in the supplementary material.

### Projections from cylindrical specimen of mouse gut

The gut is essential for life-sustaining processes, including digestion of food, absorption of nutrients and water, and excretion of waste [24]. The interactions between the enteric nervous system (ENS) and the resident immune cells are important for the normal functioning of the gut [25, 26]. In disease conditions, the spatial organisation of these cells can be disrupted affecting intercellular communication and promoting gut dysfunction. Thus, it is important to understand the cellular organisation in healthy tissue to gain insights into disease-associated changes. Light sheet fluorescence microscopy (LSFM) allows us to study the global distribution of neurons and immune cells across the different layers of the gut wall. This provides greater insight into the spatial organisation of cells in comparison to confocal images of gut wholemounts, which give larger coverage but fewer layers, or widefield images of sections, which have low coverage but include all layers [27]. We used a combination of LSFM and optical clearing [28] to acquire images of an optically cleared mouse colon. The tissue was labelled using DAPI for the nuclei, and with antibodies against Iba1 for macrophages and calcitonin gene-related peptide (CGRP) for the neuronal fibres. The challenge with this approach is the difficulty in separating the different layers of the gut within the image, and the ability to view cell types within each layer. The procedures involving mice were approved by the Monash Institute of Pharmaceutical Sciences Animal Ethics Committee (approval number 13229).

In order to study these layers in detail, spatial transformations can be applied to convert the cylindrical volume into image stacks ranging from the inside to the outside containing images which correspond to mucosa and muscularis externae layers respectively. Cartographic projections are the method of choice for visualising intensities along three-dimensional surfaces into two dimensions [29–31]. One simple form of such a projection is a maximum intensity projection along lines going from the center axis of a cylinder to its surface. Therefore, the two-channel image stack is made isotropic first, followed by a rigid transformation, which allows us to shift and rotate the volume in three-dimensional space, so as to visualize it from different perspectives. The isotropic transformed stack is subsequently processed by a cylinder transform producing an image stack where the first image corresponds to a line in the center of the colon. The images within the following slices of the stack each correspond to intensities at a given distance to that center line, going from the inner layer towards the outer layer of the gut wall. In order to get pixels at the inner surface in a two-dimensional image, the Z-position of that tissue in the cylinder stack must be determined. Therefore, we use an arg-maximum projection, also called a Z-position of maximum projection, and a Gaussian blur with a large sigma to ignore local signal intensities. After determining the Z-position of the gut wall, a corrected image stack can be determined. From this stack, we can generate two maximum projection images: One of the inner surface of the colon and one of the outer surface. As DAPI labels the nuclei of all the cells within the gut wall, the corresponding channel was used for determining the Z-position of the different layers of the gut wall. To view the macrophages within the mucosal layer (inner layer) and muscle layer (outer layer) separately, two maximum intensity projections of the corrected cylindrical projection image stacks were generated. The whole procedure is shown in Figure 4 and corresponding data and scripts are available in Supplementary Material M1.

**Figure 4:**
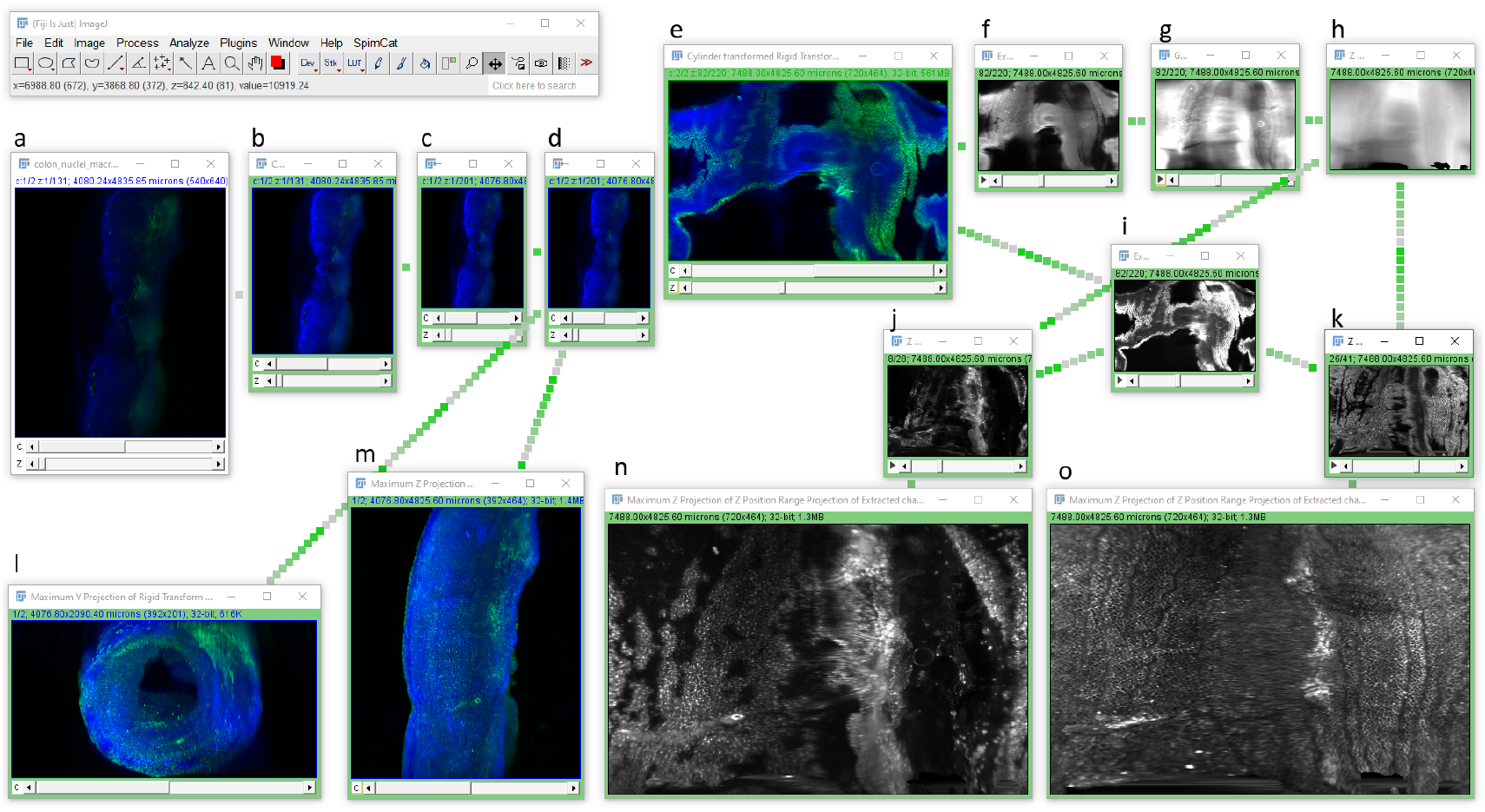
The original multi-channel image stack of a mouse colon (a) is shown labeled with DAPI in blue for the nuclei and with antibodies against Iba1 (green channel) for macrophages. The image data is pushed to the GPU (b), made isotropic (c) and rigid transformed (d). From the rigid transformed stack maximum intensity projections along y (l) and z (m) are drawn. Furthermore, the same stack is processed by a cylindrical transform (e) from which the DAPI channel is extracted (f), blurred (g) and Z-position projected (h). Together with the cylinder transfomed Iba1 channel (i), this allows users to generate corrected cylinder transformed stack of the muscularis externae of the colon (j) and mucosa (k) separately, and corresponding maximum intensity projections (n, o), respectively. The making of this IDFG is documented in Supplementary Video colon_cylindrical_projection.mp4

### Studying neighborhood relationships between cells in developing *Tribolium* embryos

Gastrulation is unarguably the major developmental event in an organism’s life. A number of 2D cell shape changes as well as 3D volumetric shape changes are seen as tissues in an embryo acquire their final shapes [32]. To understand the contribution of cell behaviors to tissue morphogenesis, it is required to provide *in toto* quantitative descriptions of cell behaviors in developing embryos. Advancements in microscopy techniques such as multi-view LSFM have enabled imaging of developing tissues with high spatial and temporal resolutions [33–35]. However, quantification of cell behaviors from such large-scale imaging data necessitates the development of fast and adaptable image analysis workflows which can be easily configured for different types of imaging data from variable biological samples and without much prior expertise in coding and programming. To demonstrate processing of such data, we acquired multi-view LSFM data set of a transgenic *Tribolium castaneum* embryo that uniformly expresses GFP in the nuclei.

An alternative to the above-mentioned strategy for cylindrical projections from a 3D volume into a 2D plane, is directly analyzing the 3D data. Therefore, GPU-accelerated image processing is beneficial to deal with the increased number of image volume elements, called voxels, which need to be processed. One basic technical challenge when analyzing such data is deriving an abstract representation of the animal’s shape. For this, it is worthwhile to use operations such as spot detection and mesh generation. In Supplementary Figure S12, an IDFG for this is set up which is available together with the used data in Supplementary Material M2. It uses background subtraction, spot detection, thresholding, binary operations, connected component labeling, label post-processing operations, mesh generation, and projections to visualize intermediate and final results.

### Digital serosa removal to study *Tribolium* embryo development

In many insect embryos the gastrulating embryonic region develops inside an outer extra-embryonic protective layer, called serosa, that forms a shell around the embryo [36, 37]. The *Tribolium* serosa shows a gradation of cell shapes and mechanical properties along the dorsal to ventral axis as the serosa shell wraps around the embryo and closes on the ventral side of the egg [38]. Once the serosa window closure is completed, the tissue stabilizes and all subsequent morphogenetic events occur in the developing embryo. Analyzing the dynamics of embryonic development have thus remained challenging. To quantify such events in *Tribolium* and potentially in other samples, we developed the IDFG shown in Figure 5 to identify a selective layer of tissue, such as the outer serosa, and digitally remove the selected layer. We remove the serosa by segmenting all nuclei and extend them virtually so that the serosal nuclei form a closed surface. Next, we use ray-tracing to identify nuclei which are selectively part of the outer surface. Deriving a binary mask, which contains all pixels except those identified as nuclei on the outer surface allows us to filter out those nuclei from the original data set. The IDFG used for this operation can be exported as script, e.g. ImageJ Macro, and modified to process all time points of the loaded data set. The generated and modified scripts are given as Supplementary Material M3. The manual addition of scale bars and timer leads to Supplementary Figure S13 showing the developing animal side by side a version of the data set where the serosa was virtually removed.

**Figure 5:**
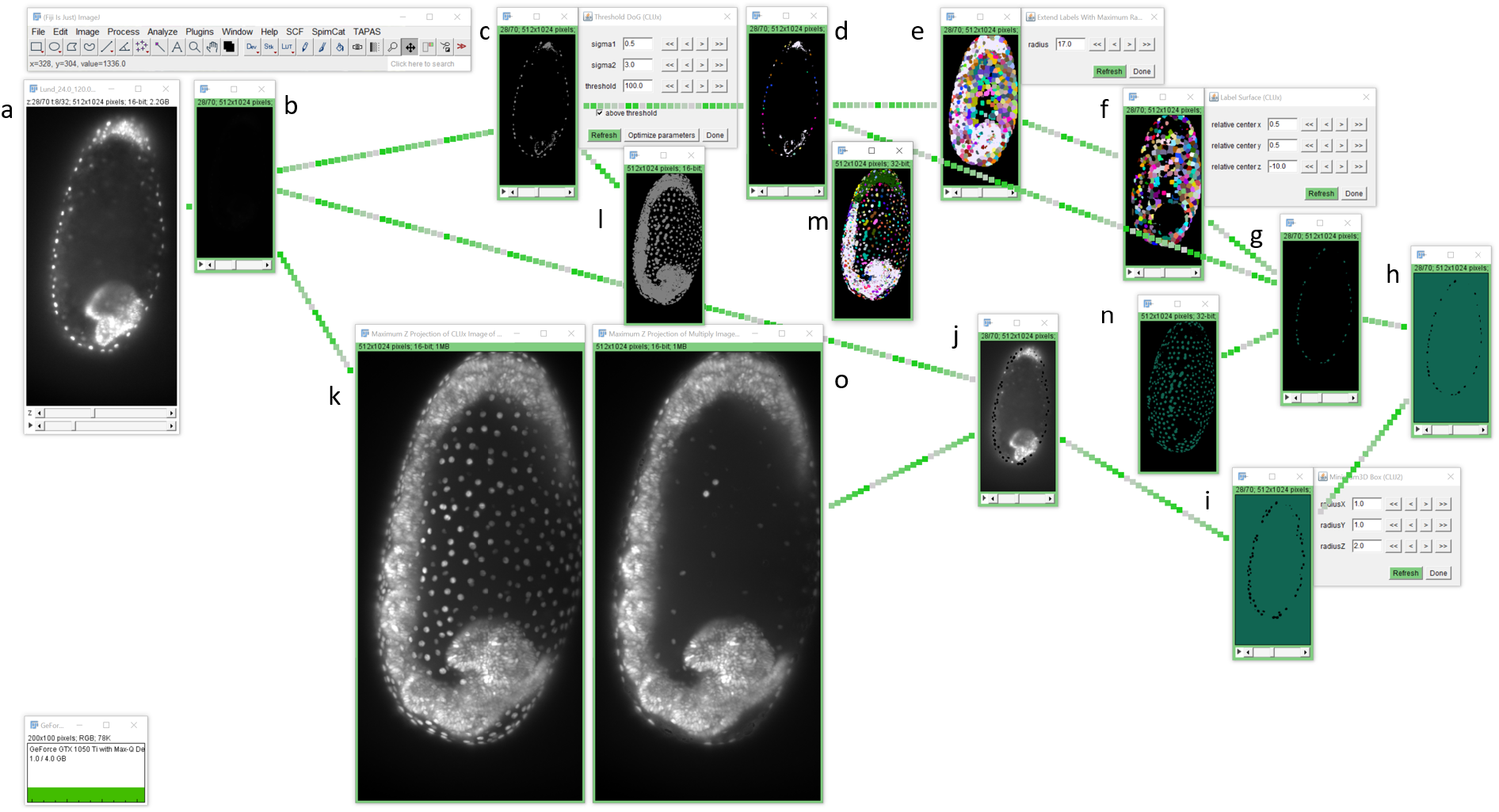
Analyzing *Tribolium* embryo development post serosa window closure: From the original 3D+time image stack (a), a 3D stack is pushed to the GPU memory (b) and thresholded (c). Then, connected component labeling is applied (e) before labels sitting on the embryo’s surface are identified (f). The thresholded image and the surface labels are combined using a binary AND operation (g) to retrieve a binary image where a binary NOT operation (h) and a binary erosion (i) are applied. The resulting binary image masks the original image (j) yielding an image stack, in which pixels of surface nuclei are set to zero. To inspect intermediate results, maximum projection of the original 3D stack (k), thresholding (l), labeling (m), binary surface nuclei (n) and the embryo data set with removed surface nuclei (o) are shown. See also Supplementary Video tribolium_surface_removal.mp4

### Cell classification on large 2D histological mouse brain sections

Cancer treatment applies various approaches to cure or delay tumor growth. For example, radiotherapy induces DNA damage, which inactivates dividing tumor cells and can induce cell death. Unfortunately, treatment may also affect surrounding normal tissue, leading to potential long-term side effects in surviving patients. Suitable preclinical models are crucial to investigate these side effects, find potential predictors, and test new treatment approaches. A recently established mouse model to study radiation injury in brain tissue [39] uses γH2AX, a protein formed during the DNA doublestrand repair mechanism, to visualize immediate effects. Costaining with DAPI, a marker for cell nuclei, offers the possibility to investigate the damaged cell fraction and correlate the result with the applied radiation dose. Mouse irradiation experiments were performed respecting European (EU Directive 2010/63/EU) and national animal welfare guidelines under approval number 24.1-5131/394/50 (Landesdirektion Sachsen).

For image analysis of large 2D histological specimen it is recommended to be performed tile-by-tile by extracting a smaller part from the large 2D image and splitting the channels. Afterwards, segmentation is applied to both channels individually. Then, counts can be determined per channel and tile to estimate DNA damage [39]. Alternatively, overlap measurements such as the Jaccard Index can be used to derive a per-cell probability of detected DNA-damage. This approach, visualized as IDFG in Figure 6, allows an instant comparison of different regions of the tissue. The corresponding data and scripts are available in Supplementary Material M1. After exporting the IDFG as script, the procedure can potentially be executed on the whole brain slice, in order to derive a map of DNA damage. Therefore, additional coding efforts are required.

**Figure 6:**
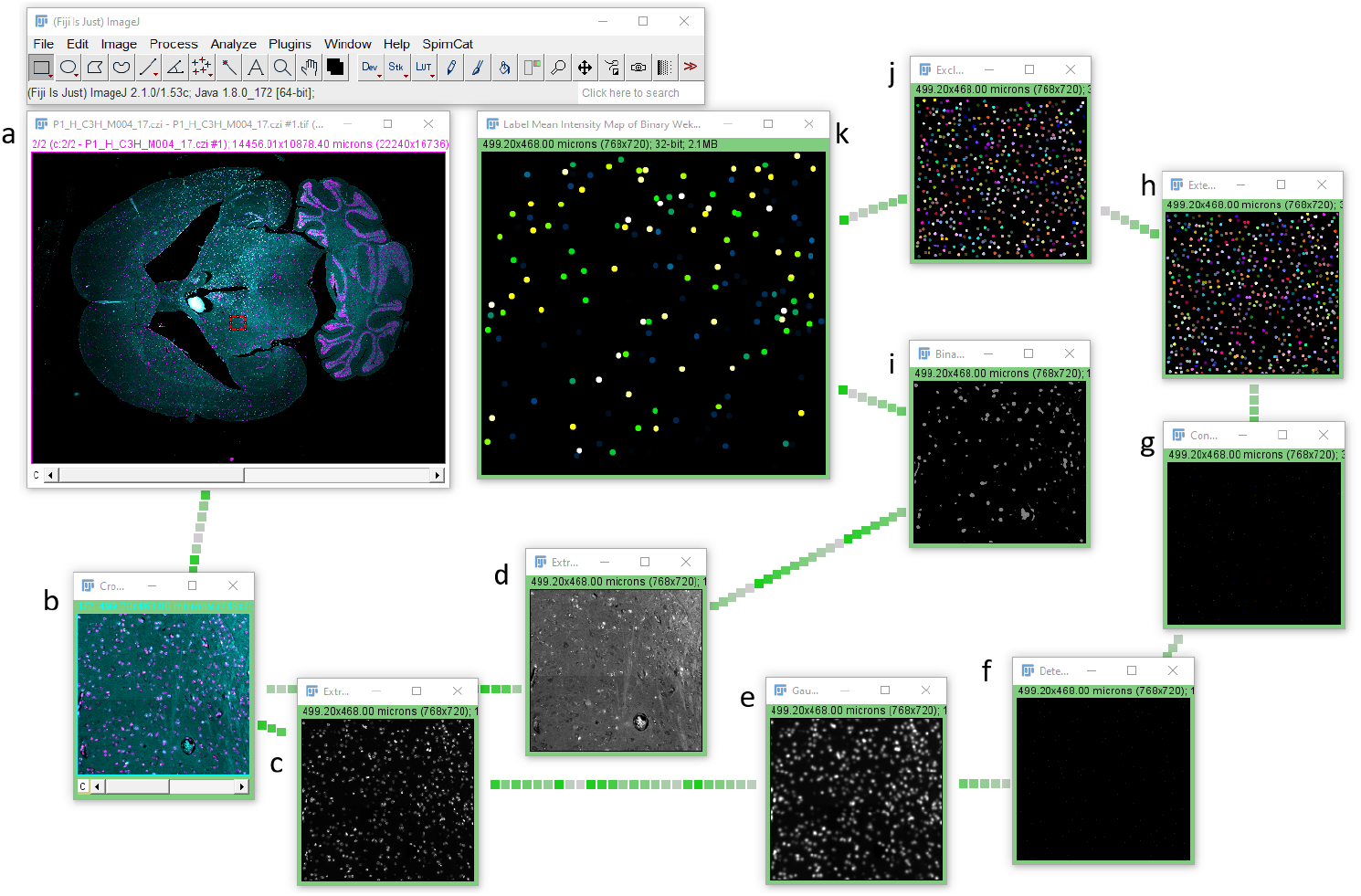
Exploring a multi-channel mouse brain slice to determine DNA damage, from the original 2D image (a), a regions is cropped out (b) and the two channels are extracted (c, d). Starting from the DAPI channel (magenta), Gaussian blur (e), spot detection (f), connected components labeling (g), label extension (h) and exclusion of labels at the image border (j) are applied. The *γ*H2AX channel (cyan) is binarized using a binary Weka pixel classifier. The label map and the binary image are then used to compute the amount of overlap between individual nuclei labels and the binary image. This measure can be interpreted as surrogate parameter for DNA damage. Such an IDFG allows to explore a large 2D slice tile-by-tile, as shown in Supplementary Video mouse_brain_cell_classification_dna_damage.mp4

## 4 Discussion

We have presented an overview of CLIJ-assistant, a userfriendly tool for designing GPU-accelerated image analysis workflows in ImageJ. Across multiple sample types and imaging technologies, our approach enables visualization and quantification of tissue and cellular behaviors with an interactive and configurable toolkit. The necessary user actions towards a well-suited image analysis workflow are facilitated using automatic suggestions by an expert system, pre-selection of suitable subsequent operations and automatic parameter optimization. While workflow design does not involve programming skills, it is possible to export workflows in various programming languages and to deploy them to other platforms. This approach bridges communities such as biologists and computer scientists, and users of different image analysis platforms.

The presented user-interface for GPU-accelerated IDFG design has one major limitation: GPU memory. It limits the number of image stacks which can be handled on screen - which is limited by screen size anyway. This suggests to think about necessary number of processing steps: The smaller the number of processing steps, the easier the justification of a workflow. Furthermore, the built-in Fiji-plugin generator allows organizing memory consumption of workflows: If the users employ a chain of *n* operations in multiple projects blocking *n* times image size in GPU memory, they can generate a Fiji plugin from this chain. When this plugin is introduced in future workflows, it spares *n* − 1 times image size compared to earlier workflows. In that way, users learn how to structure workflows into sub-routines and how to reuse these sub-routines in larger workflows. For processing images larger than available GPU-memory, tile-by-tile processing strategies similar to the demonstrated mouse-brain cell classification workflow need to be developed. These strategies can be straight-forward: for example simple image filtering can be done tile-by-tile with overlapping margins and a result image can be assembled right away. If operations such as watershed, skeletonization or connected component analysis are part of the workflow, tiled processing is not trivial and thus, efficient GPU-accelerated strategies for this will part of our future work.

The used CLIJ plugin system is a mixture of the ImageJ2 [9] and ImageJ [3] extension systems, including conceptual ideas from the Insight ToolKit (ITK) project [40]. First of all, plugin-discovery is driven by ImageJ2’s plugin system. With this mechanism custom plugins on the Java class path can be discovered and offered to the user in right-click menus, the search bar and in the auto-completion of the script editor. Secondly, ImageJ’s Macro Extensions are used to drive the macro-compatibility. The underlying architecture suggests using arrays of objects as parameter lists. This approach was chosen because it paves the path towards compatibility with QuPath and Icy. Furthermore, passing arrays of objects allows passing of one image over to two CLIJ plugins: The first plugin receives the image as output image; the second plugin receives it as input. In that way, plugins can be chained together in a fashion similar to the mechanism used in ITK. Thus, CLIJ operations have a state. They are not just functions that are executed once. They are parameterized functional objects. This allows repeated execution of operations without the need for allocating and freeing memory repeatedly. One potential disadvantage of the approach is the limited number of supported types of parameters: Only numbers, strings, arrays and images are supported. On the one hand this may be perceived to be limiting by software developers. On the other hand it simplifies image processing for the end-user.

Another aspect of interoperability is the frontier between Java, Python and C++ based image processing libraries. Also other image processing libraries, such as SimpleITK, support all three developer communities by providing access to its functionality in these three languages. We are about to extend CLIJ towards Python and C++ as demonstrated above in the clEsperanto project to make it available to a broader audience. Therefore, the re-implementation of filters for the Python and C++ side might appear like a lot of effort. However, there are substantial benefits to the community: Programmers used to ImageJ’s scripting languages can copy over code snippets to Python driven Jupyter notebooks, and Python programmers can provide code snippets from their workflows to users of ImageJ, Icy or QuPath. This approach bridges communities and facilitates knowledge exchange. Most importantly, it allows workflow designers to focus on the scientific question rather than technical implementation details of analysis procedures. Furthermore, as increased efforts are required for automated testing and deployment of the Python, Java and C++ environments of clEsperanto, the project has the chance to achieve high quality implementations of image processing operations.

The user-interface of the CLIJ-assistant is under development, and this preprint serves as a base for discussion between the user and developer communities. Thus, the interface may change in the future. We want to highlight that if users generate code using CLIJ-assistant employing CLIJ2 operations to process their images in ImageJ, Fiji, Icy or Matlab, these workflows will continue to work even if the CLIJ-assistant changes. This is also true for generated Java-based Fiji-plugins as long as they are limited to CLIJ2 operations. A strong delimitation between CLIJ2 operations and the presented IDFG user interface ensures that. CLIJx operations may also be subject to change, as the x in CLIJx stands for ‘experimental’. Nevertheless we would emphasize that user feedback on these operations is guiding us towards the next generation of CLIJ and allows us to make decisions on which CLIJx functionality should be included in future releases.

## 5 Conclusions

The presented CLIJ-assistant enables interweaving of common image analysis operations for comprehensive workflow construction. Interactive design of image data flow graphs with instant feedback through GPU-accelerated image processing enables new ways of intuitive learning and teaching image analysis in life-sciences and beyond. Therefore, we expect a swift adaptation by many research fields beyond the examples shown here. Further, using the suggested concept of IDFG design enables sharing image processing routines between experts in a programming language-independent fashion. Users can export code for multiple platforms. This compatibility enables the exploration of other platforms in individual projects and, thus, bridges communities with the aim to solve the underlying analysis questions rather than platform-specific implementation related technical challenges.

## Acknowledgements

We would like to thank everybody who helped motivating, developing and testing CLIJ-assistant and the colleagues who supported us in performing the experiments of which we re-used data for demonstrating the potential of the CLIJ-assistant. In particular thanks go to Armin Lühr (TU Dortmund), Barbara Treutlein (D-BSSE, ETH-Zürich), Benjamin Pavie (VIB Leuven), Bert Nitzsche (PoL TU Dresden), Bradley Lowekamp (NIAID Washington), Bram van den Broek (Netherlands Cancer Institute Amsterdam), Bruno C. Vellutini (MPI CBG Dresden), Christian Tischer (EMBL Heidelberg), Cläre von Neubeck (University Hospital Essen), Curtis Rueden (University of Madison), Deborah Schmidt (CSBD/MPI CBG Dresden), Elke Beyreuther (OncoRay Dresden), Florian Jug (CSBD/MPI CBG Dresden), Gayathri Nadar (MPI CBG Dresden), Irene Seijo Barandiaran (MPI CBG Dresden), Johannes Girstmair (MPI CBG Dresden), Johannes Müller (OncoRay Dresden), Kisha Sivanathan (Harvard Medical School Boston), Liane Stolz-Kieslich (DKTK Dresden), Malte Gotz (OncoRay Dresden), Marion Louveaux (Institut Pasteur Paris), Matthias Arzt (CSBD/MPI CBG Dresden), Mechthild Krause (OncoRay, University Hospital Carl Gustav Carus, Dresden), Noreen Walker (MPI CBG Dresden), Pete Bankhead (University of Edinburgh), Romain Guiet (EPFL Lausanne), Sebastian Munck (VIB Leuven), Stein Rørvik, Stéphane Dallongeville (Institut Pasteur) and Tanner Fadero (U Chapel Hill) Furthermore, the constant support by the Image Science and the NEUBIAS communities is acknowledged.

The imaging in this work was supported by the Light Microscopy Facility of the CMCB Technology Platform at TU Dresden and the Advanced Imaging Facility at MPI-CBG Dresden.

R.H. was supported by the German Federal Ministry of Research and Education (BMBF) under the code 031L0044 (Sysbio II) and by the Deutsche Forschungsgemeinschaft (DFG, German Research Foundation) under Germany’s Excellence Strategy – EXC2068 - Cluster of Excellence Physics of Life of TU Dresden.

## Authors contributions

RH, DV, PR and TS acquired the *Tribolium* and mouse image data. RH, SR, TJL, JNI and PR programmed the software. PR, DV, AJ and TS tested the software and guided development towards usability and applicability. DPP, PT and EWM supervised the project. RH, AJ, DV, PR and TS wrote the manuscript with input from all co-authors.

## Competing Interests

Research in the laboratories of DPP is funded in part by Takeda Pharmaceuticals International.

## Code availability

The open source code, documentation and installation instructions are available online http://clij.github.io/assistant/.

## Supplementary material

Scripts and videos are available online https://doi.org/10.5281/zenodo.4276076

## A Supplementary material

**Figure S1:**
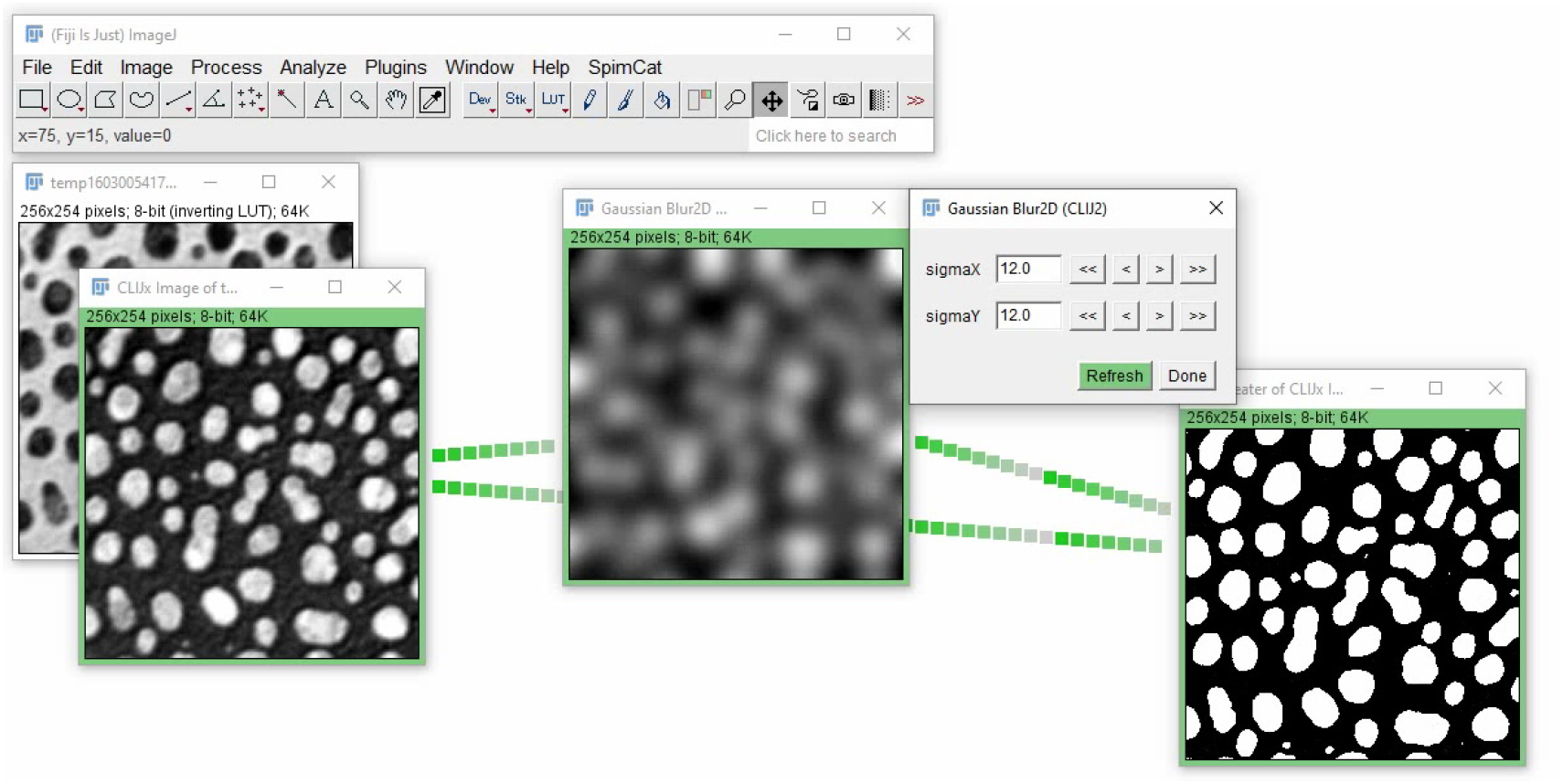
Basic IDFG assembly involves setting up connected image processing operations step by step. In contrast to classic ImageJ, all configuration dialogs can stay open to change parameters and the resulting impact on the graph can be observed. In this example, the blobs.gif ImageJ example image is pushed to GPU and smoothed using a Gaussian blur. Afterwards, a binarization is applied. The procedure is shown in detail in Supplementary Video basic_usage.mp4

**Figure S2:**
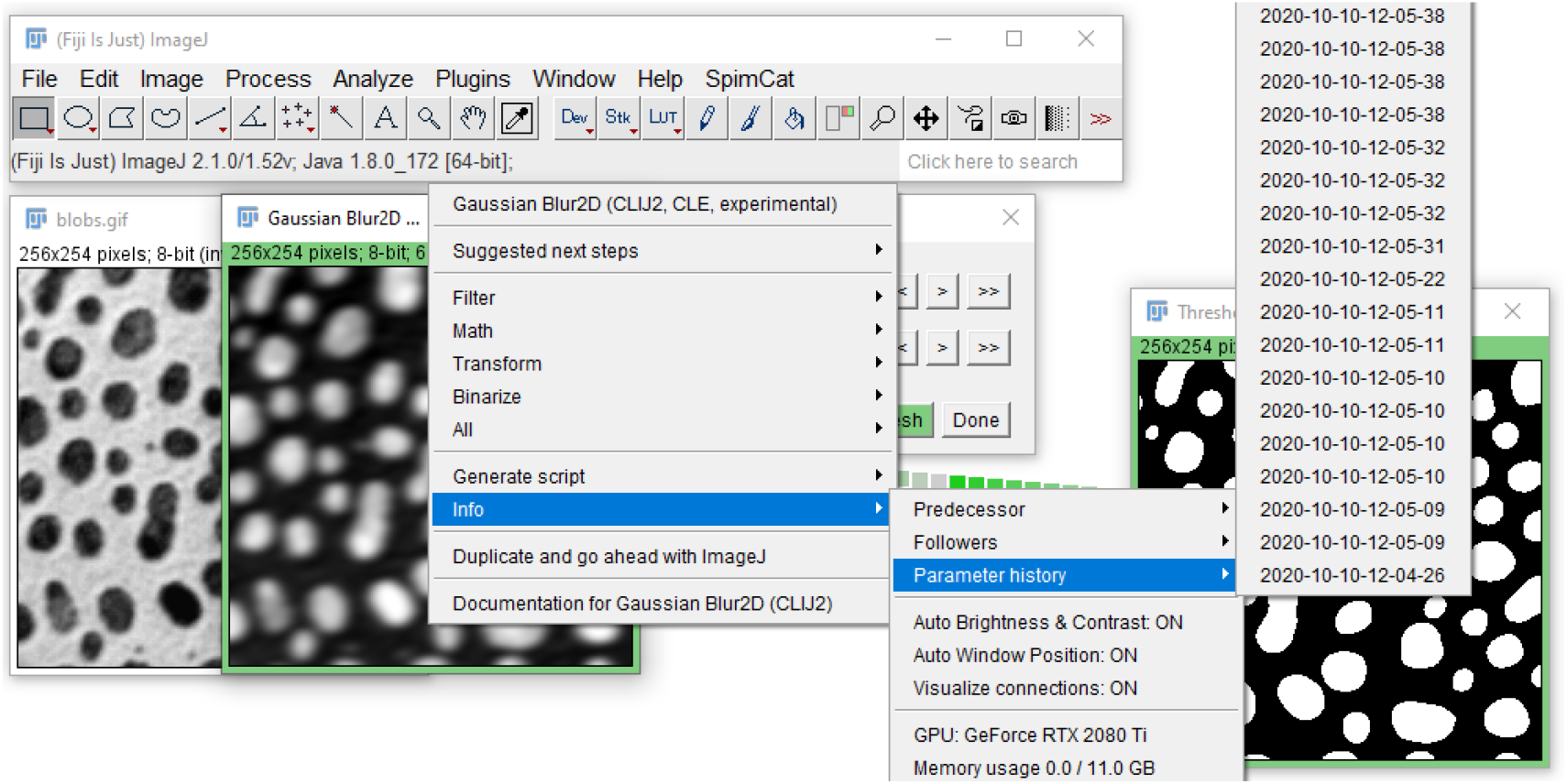
Before every parameter change, all parameters are stored internally in the parameter history. Later on, the user can go back to a parameter configuration at arbitrary time points from the shown right-click menu.

**Figure S3:**
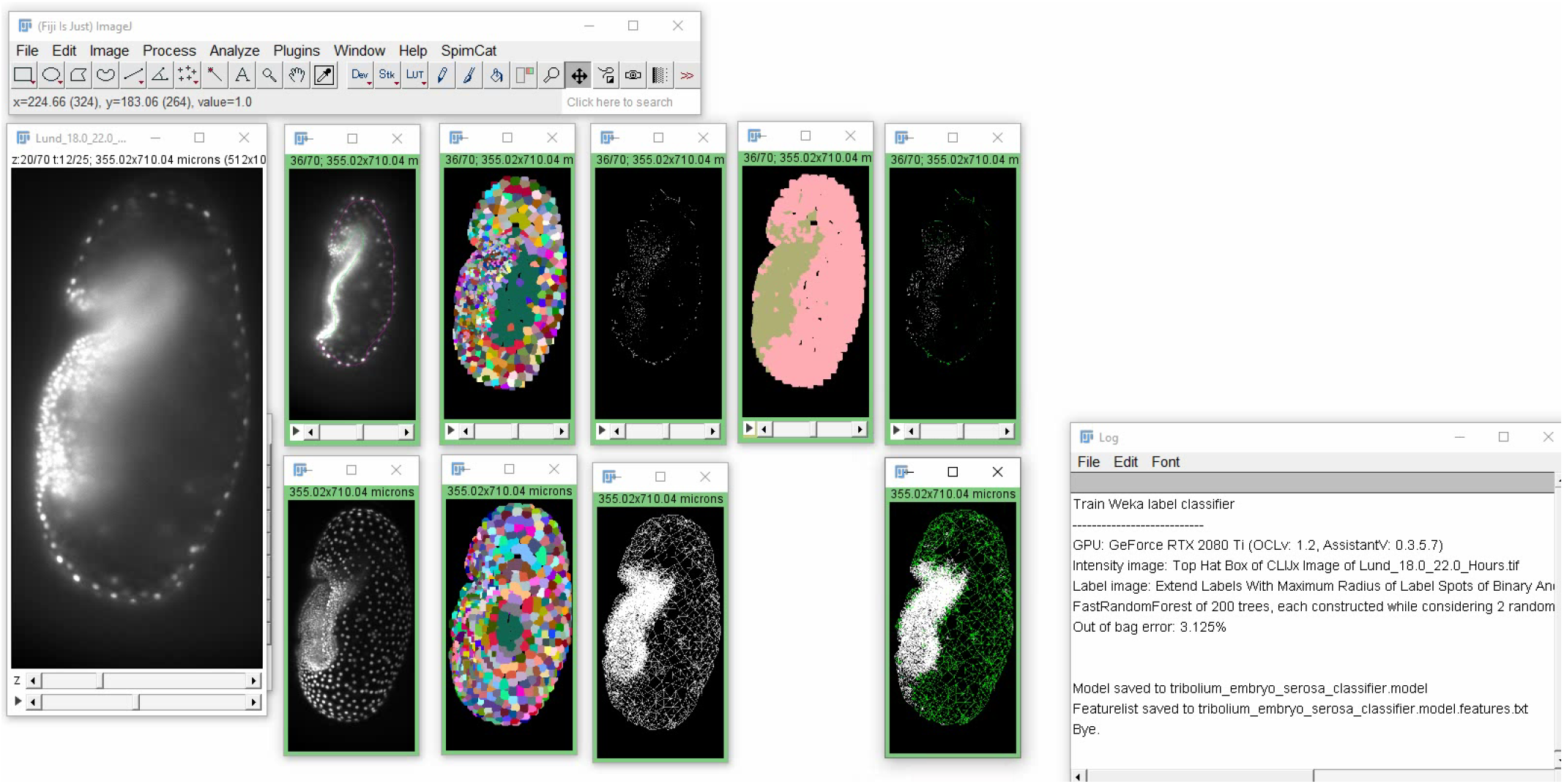
Weka based label classification can be incorporated into IDFGs, for example to differentiate embryo and serosa in *Tribolium castaneum*. The random forest takes the background subtracted image stack (Second column), a corresponding label image stack (third column) and pre-defined annotation to train a model to differentiate embryo and serosa based on intensity and topological properties of segmented objects. By multiplying a neighbor mesh (fourth column) with predicted classification image (fifth column), a classified mesh can be generated (sixth column). See also Supplementary Video weka_label_classifier_short.mp4

**Figure S4:**
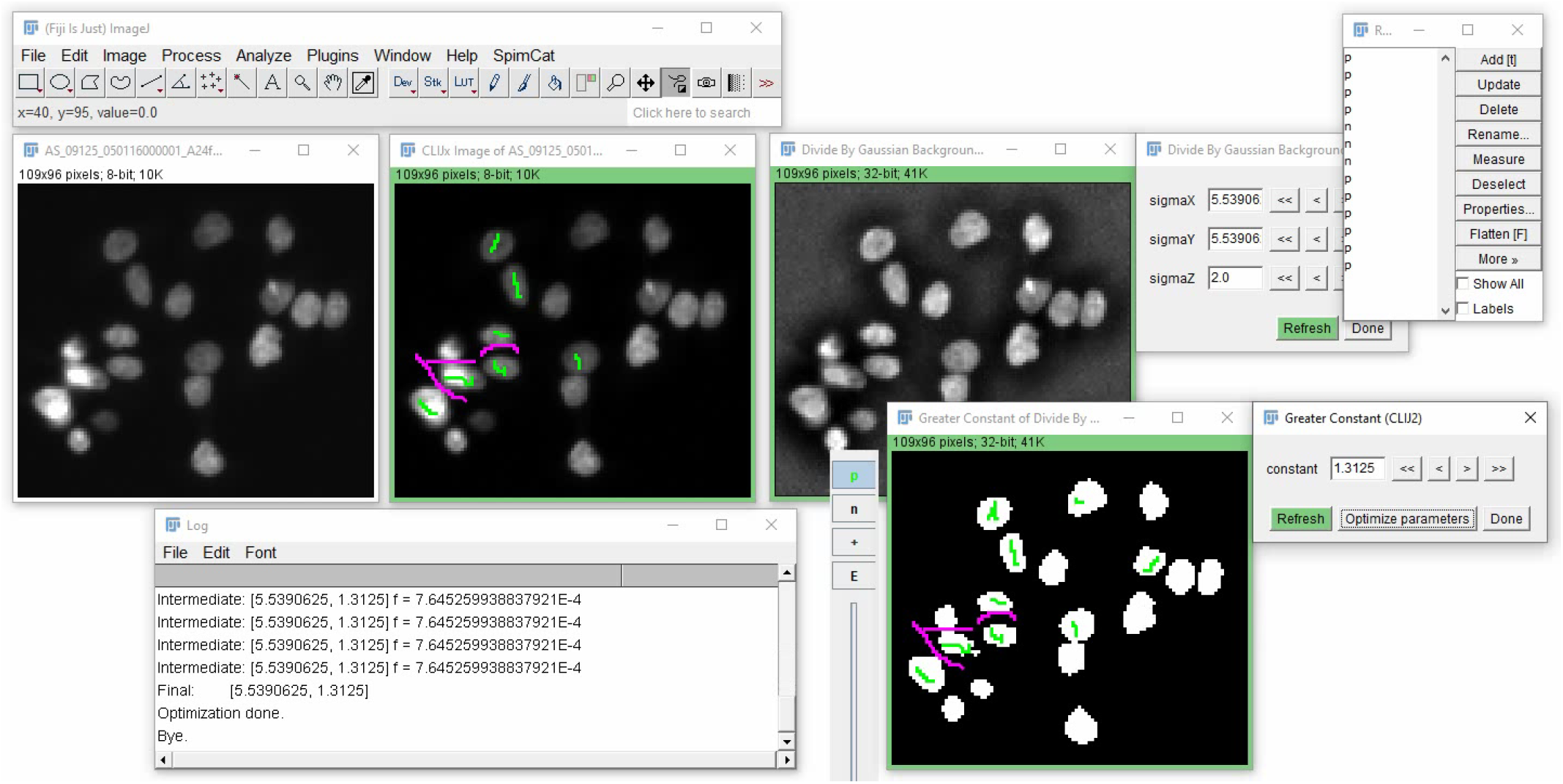
After annotation of an input image, parameter optimization can be used to configure the graph to minimize the error in a given binary result image. The processed image data was taken from the Broad BioImage Challenge [41]. See also: optimize.mp4

**Figure S5:**
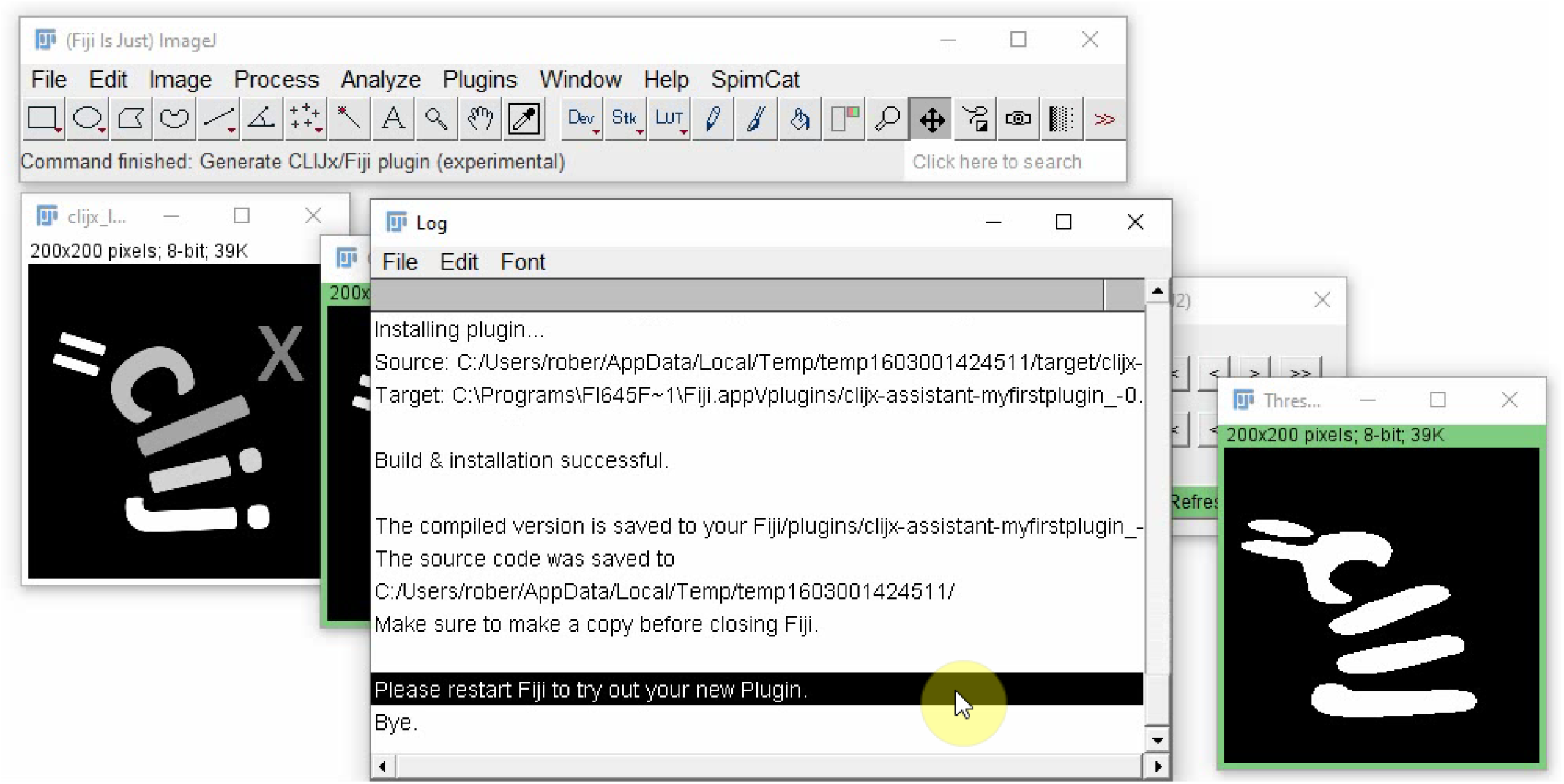
IDFGs can be converted into Fiji plugins. For this, Java code is generated, inserted into a Maven based CLIJx/Fiji plugin template and compiled. The user could also open the generated code in an integrated development environment and refine it. The fully automatic compilation and installation procedure is shown in Supplementary Video fiji_plugin_generator.mp4

**Figure S6:**
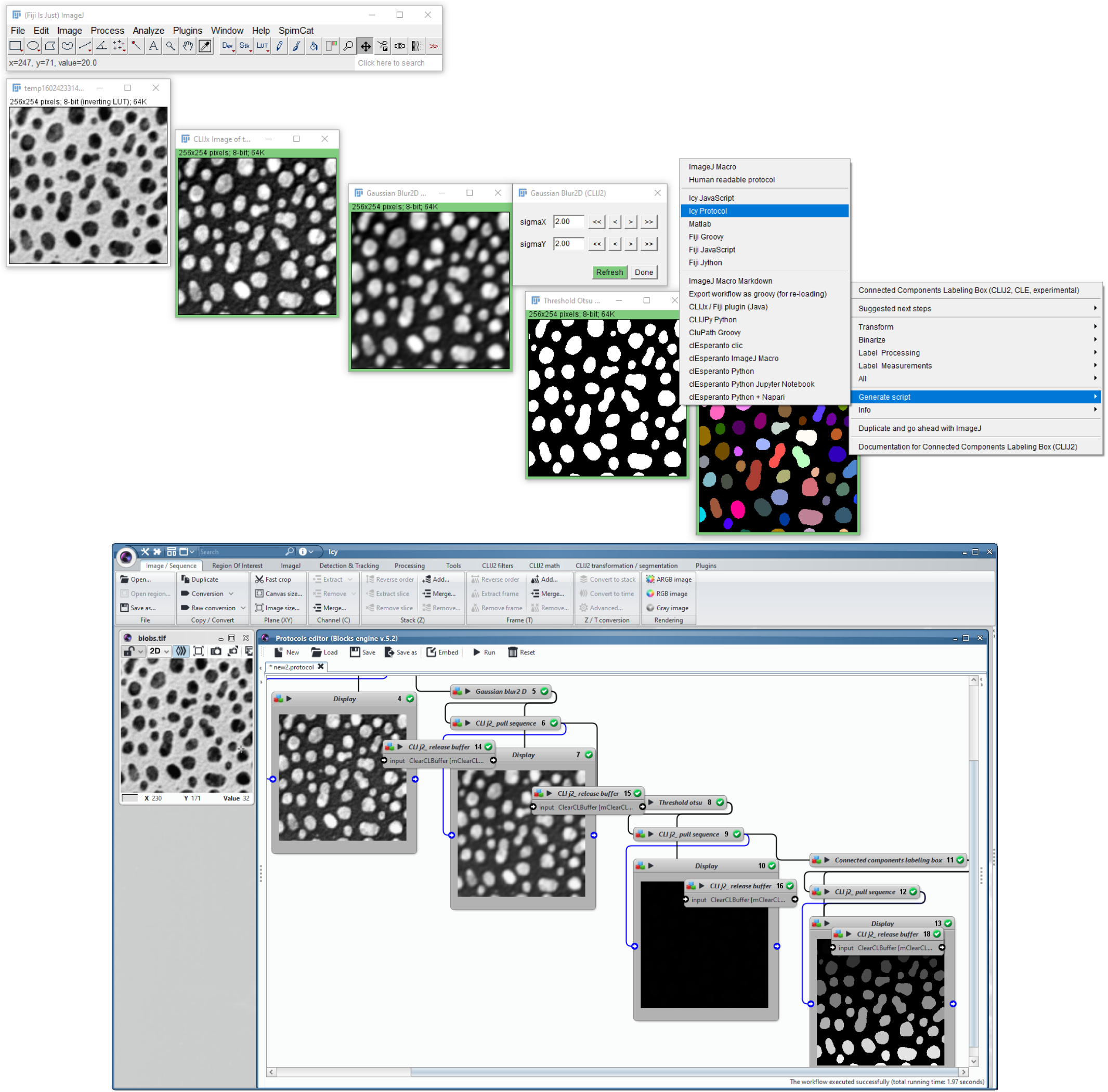
IDFGs can be exported as Icy protocols via the right-click menu, as shown in the top row. If Icy is configured correctly in ImageJ, it opens the Icy-protocol, which corresponds to the IDFG immediately. The procedure is demonstrated in blobs_icy_export.mp4

**Figure S7:**
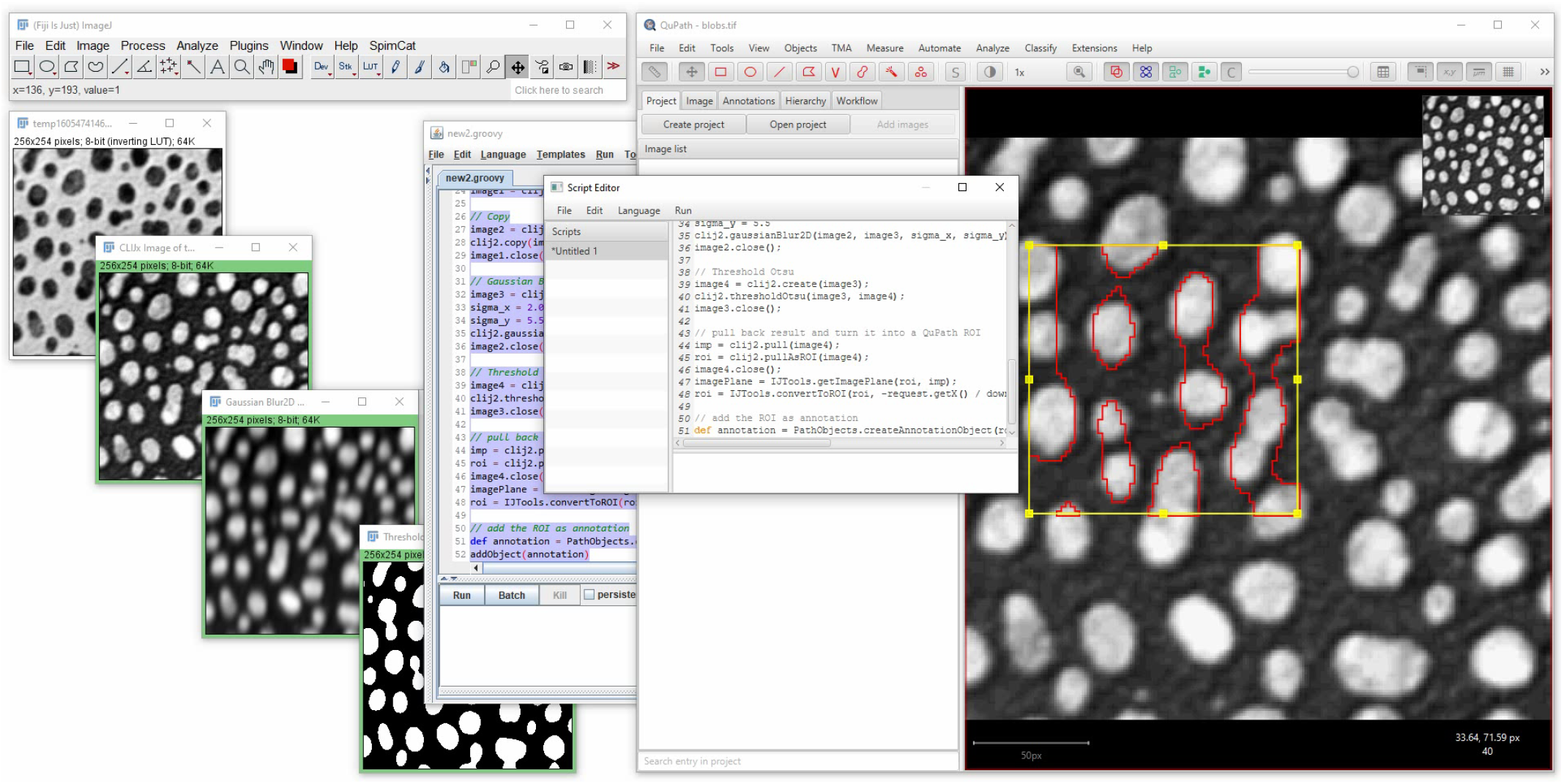
IDFGs which result in two dimensional binary images can be exported as QuPath compatible Groovy script to produce regions of interest in QuPath as demonstrated in clupath_script_export.mp4

**Figure S8:**
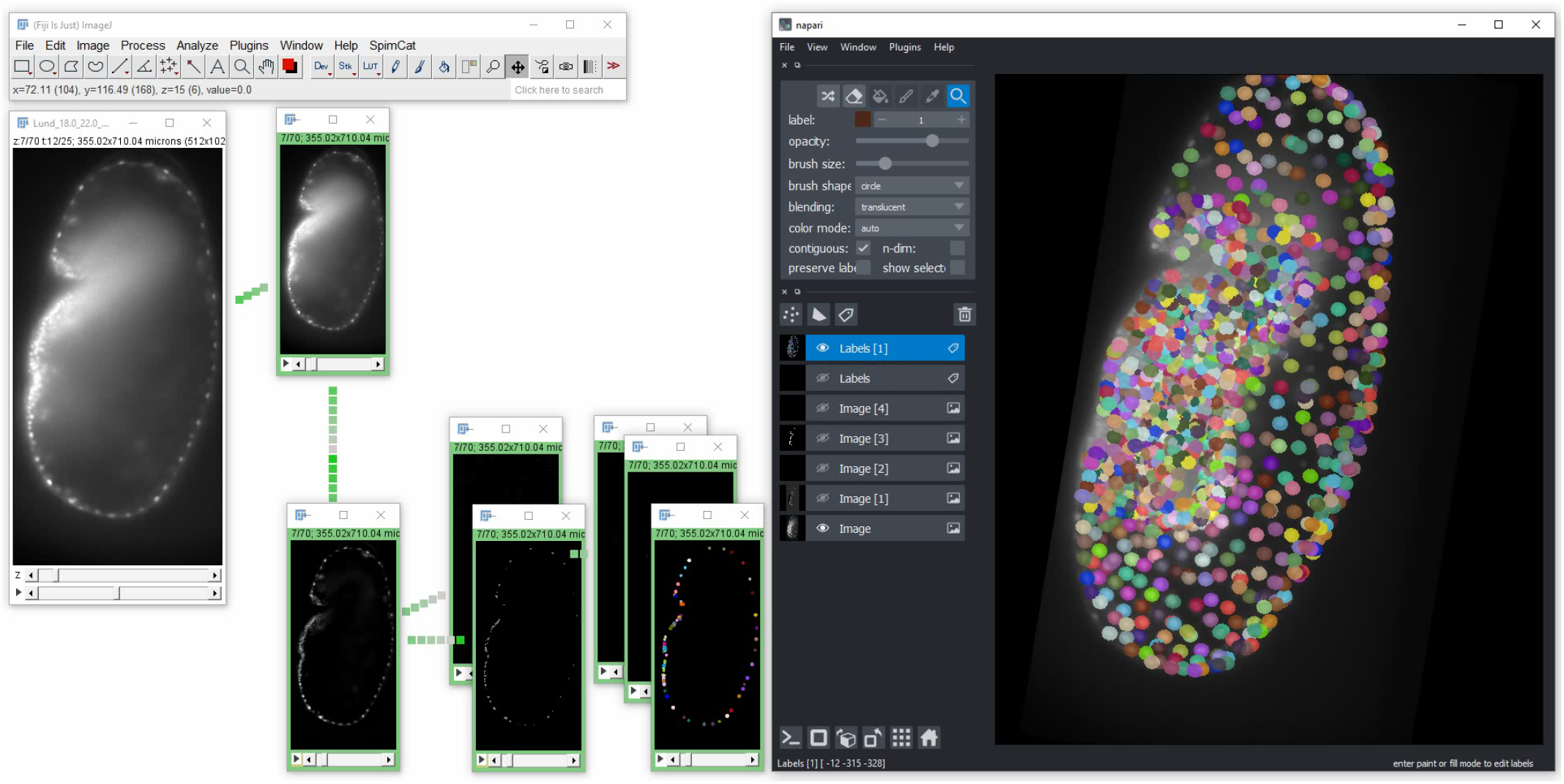
IDFGs can be exported as a Python script, which uses napari to visualise image processing results of intermediate steps as layers, as shown in the center. Napari can be opened from configured IDFGs as shown in Supplementary Video tribolium_spot_detect_napari.mp4

**Figure S9:**
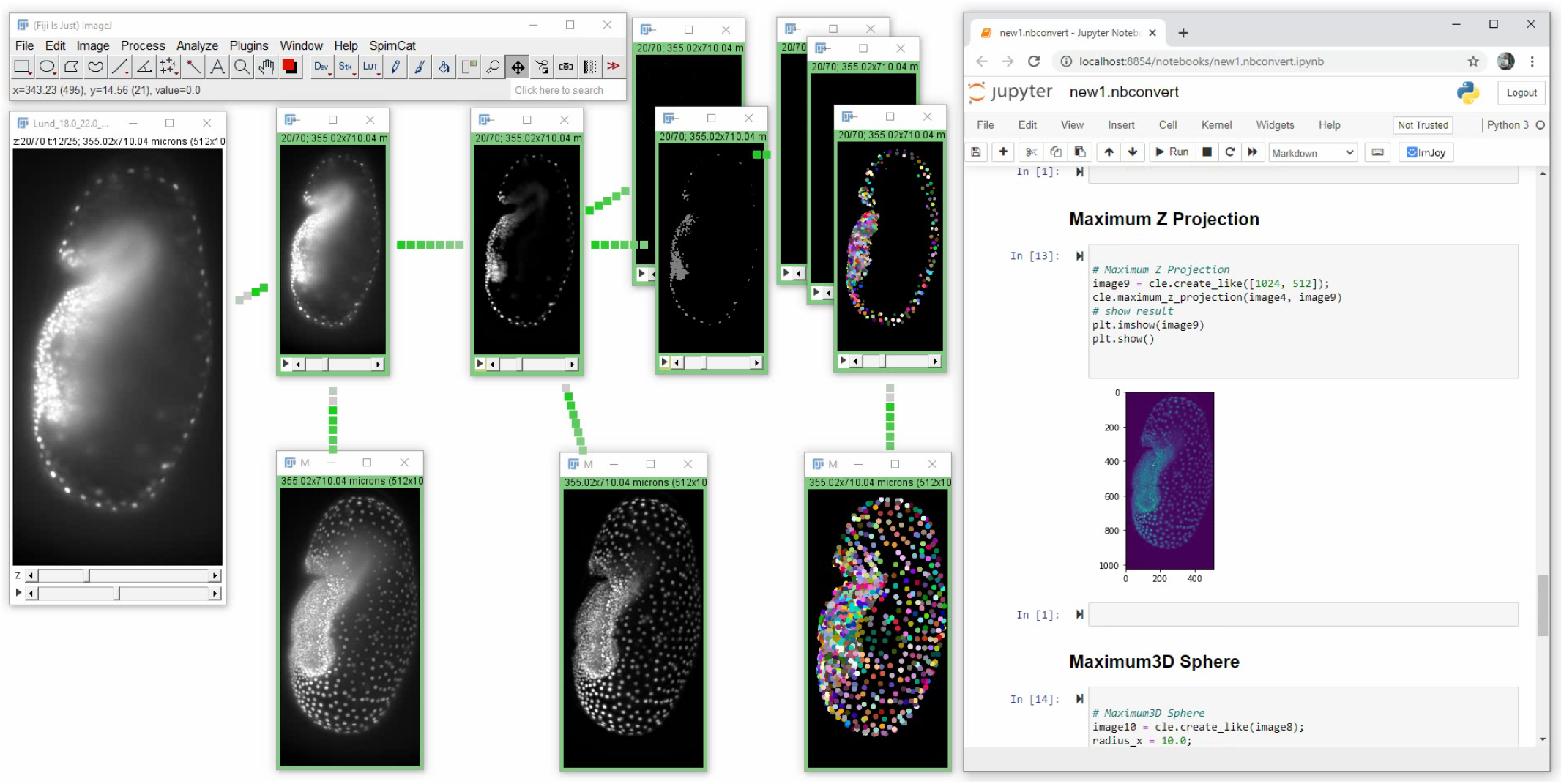
IDFGs can be exported as Python 3 Jupyter notebook as shown in the center which contains the image processing steps as code cells. See also Supplementary Video tribolium_spot_detect_max_proj_jupyter.mp4

**Figure S10:**
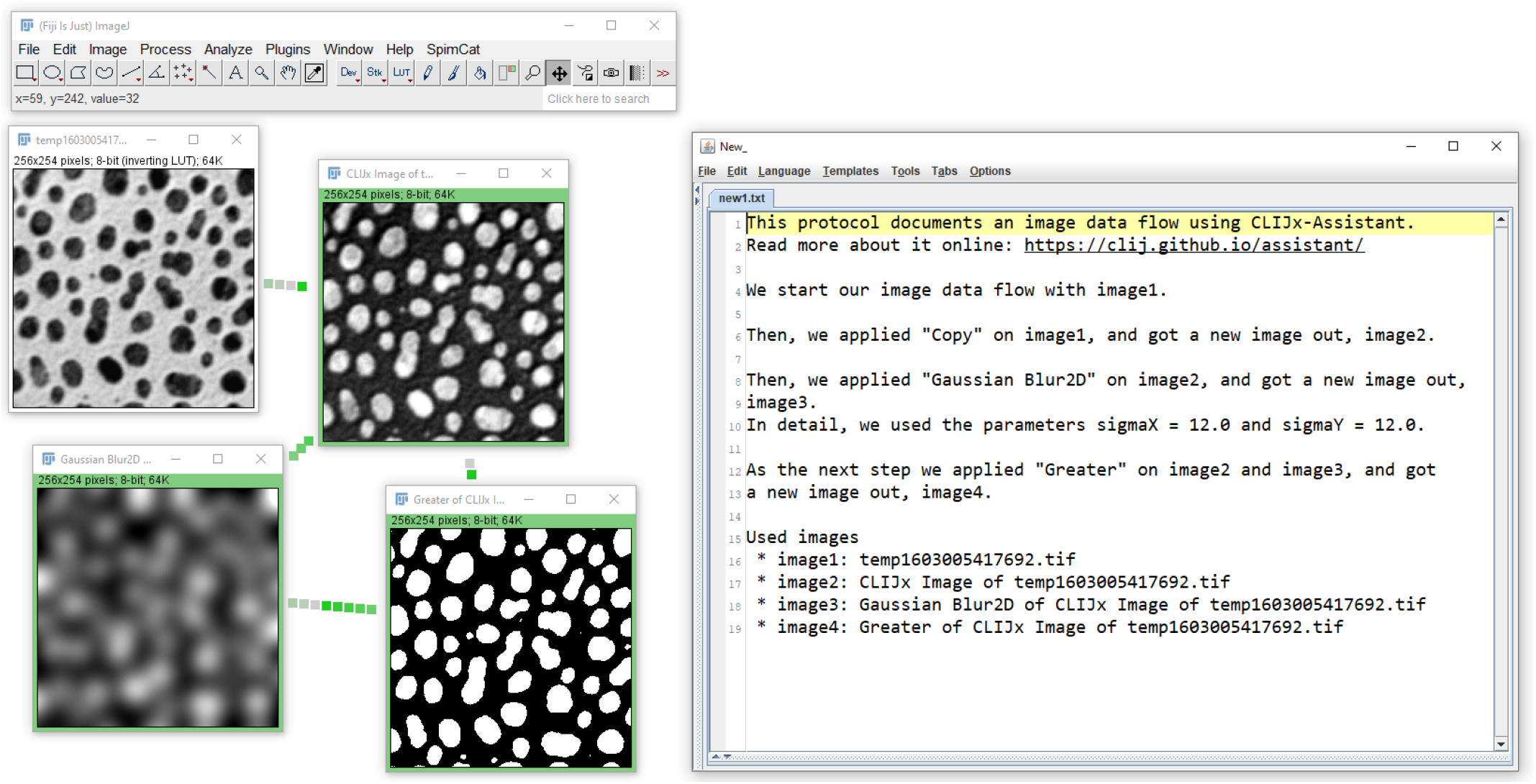
A human readable protocol generator turns IDFGs (left) into documentation text (right).

**Figure S11:**
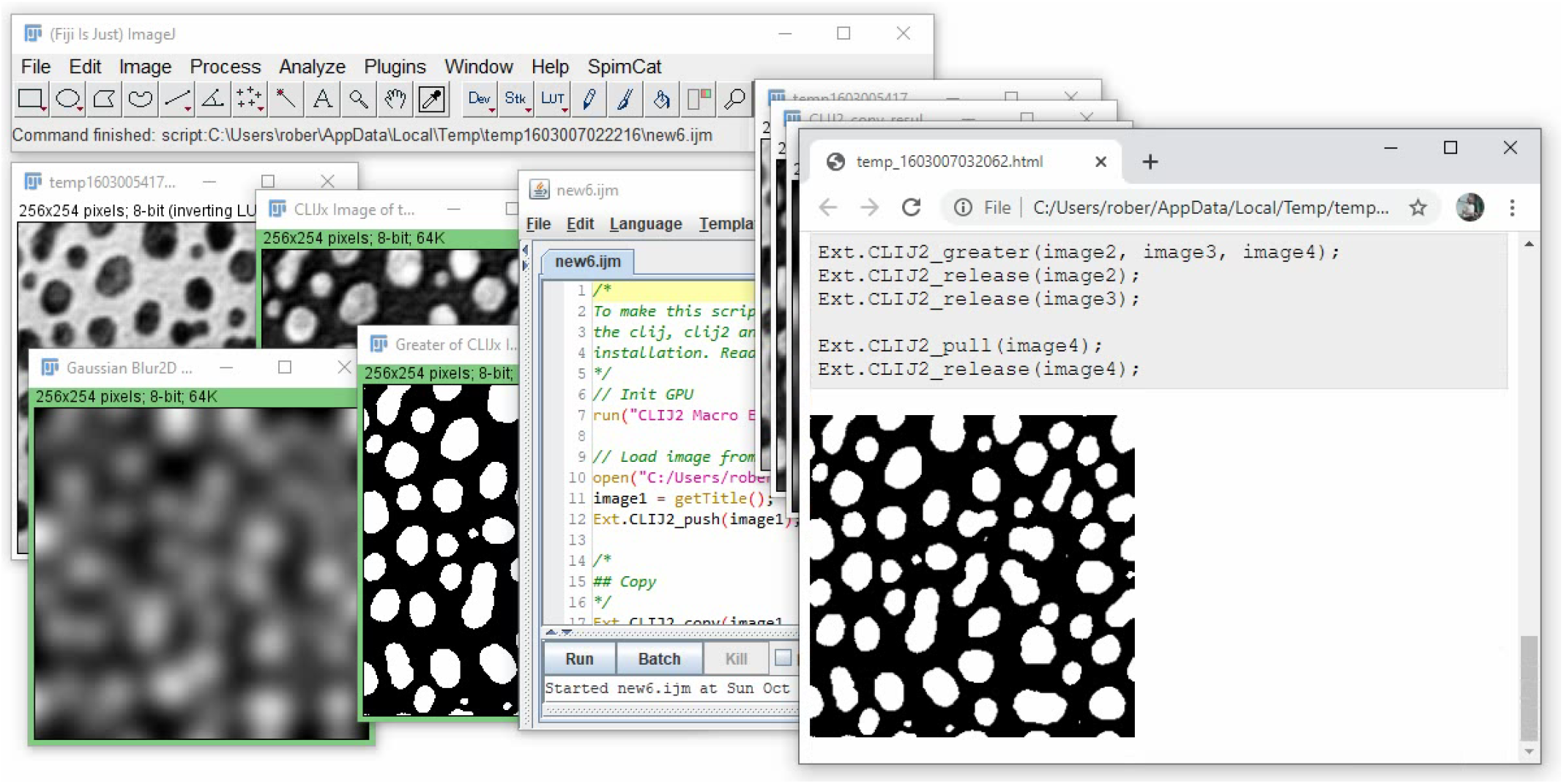
An IDFG (left) can be exported as an ImageJ Macro Markdown notebook (right), which is a hybrid between an executable script and a human readable documentation of the workflow in a notebook fashion. See also Supplementary Video ijmmd.mp4

**Figure S12:**
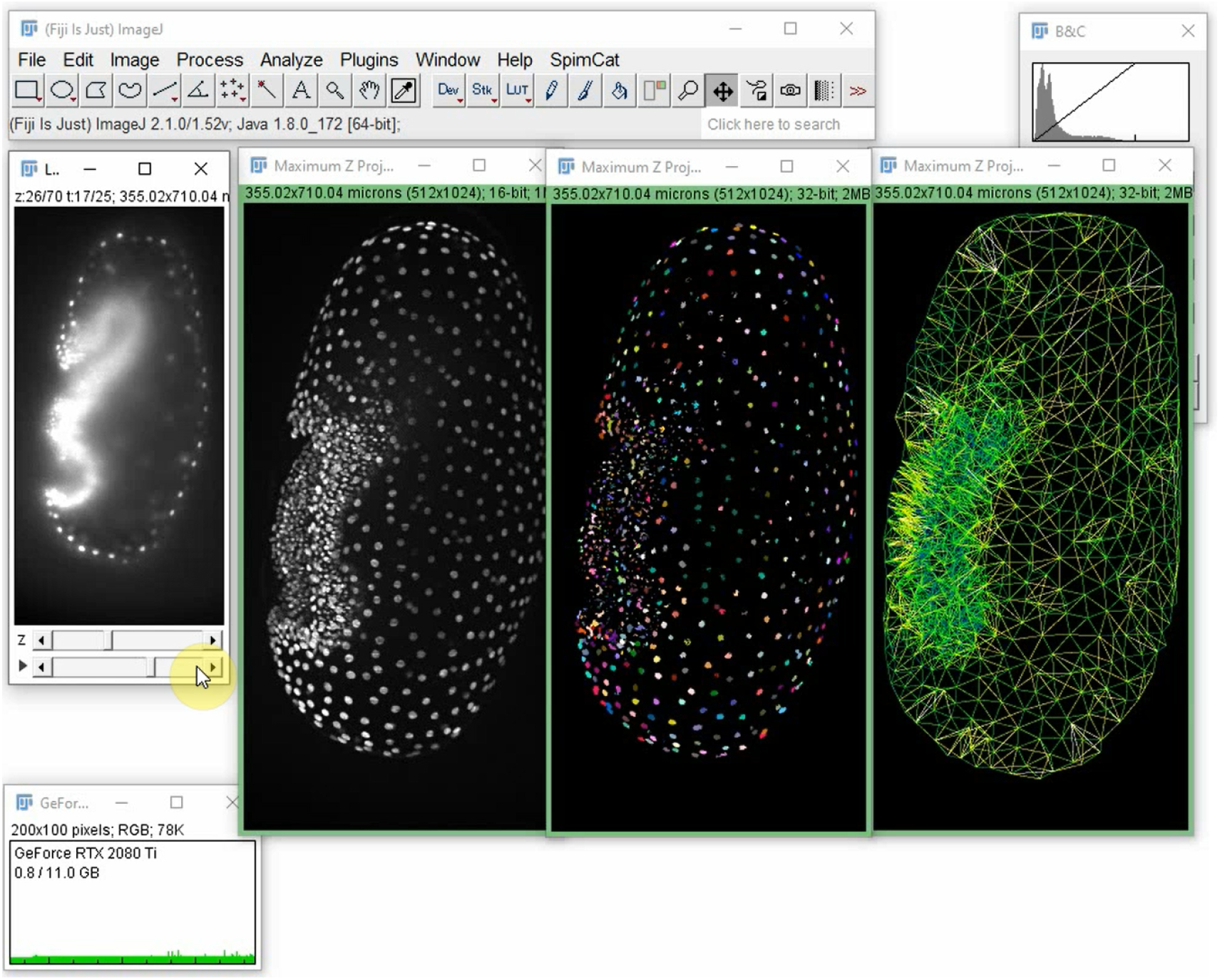
Setting up an IDFG which takes raw light sheet microscopy time lapse image stacks as input and derives meshes connecting neighboring nuclei is shown in Supplementary Video tribolium_cell_neighborhood_distance_visualization.mp4

**Figure S13:**
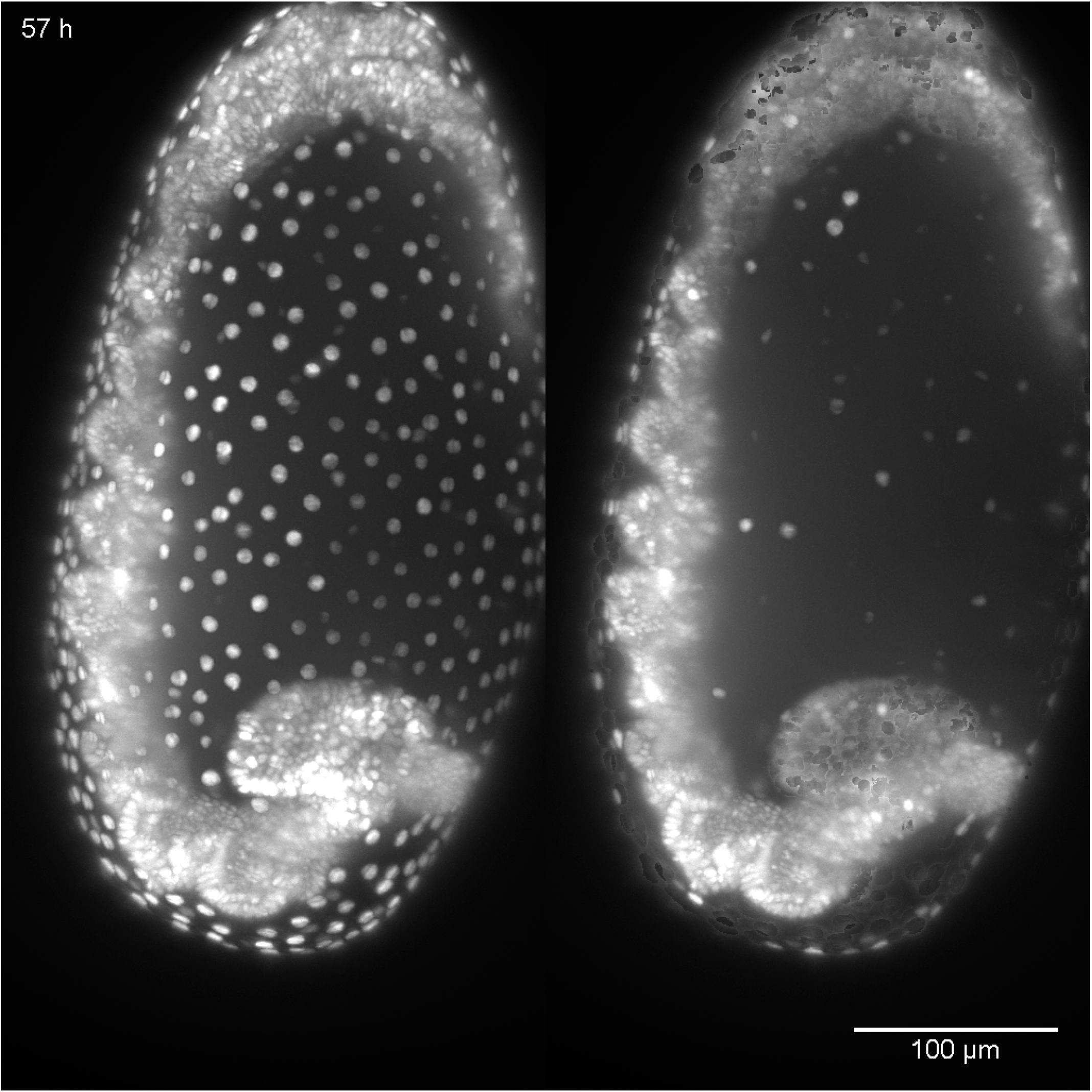
By removing the outer layer of detected nuclei, presumably serosa, improved visualization of *Tribolium* embryo development is achieved. See also video: tribolium_surface_removal_result_video.gif

**Figure M1:**
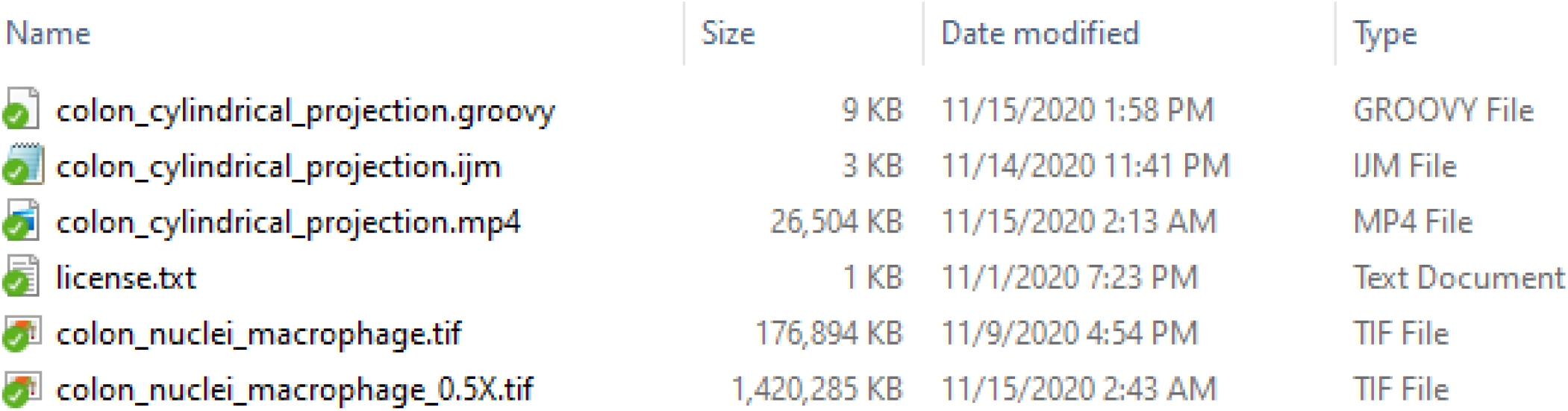
Supplementary material: Mouse colon data + scripts

**Figure M2:**
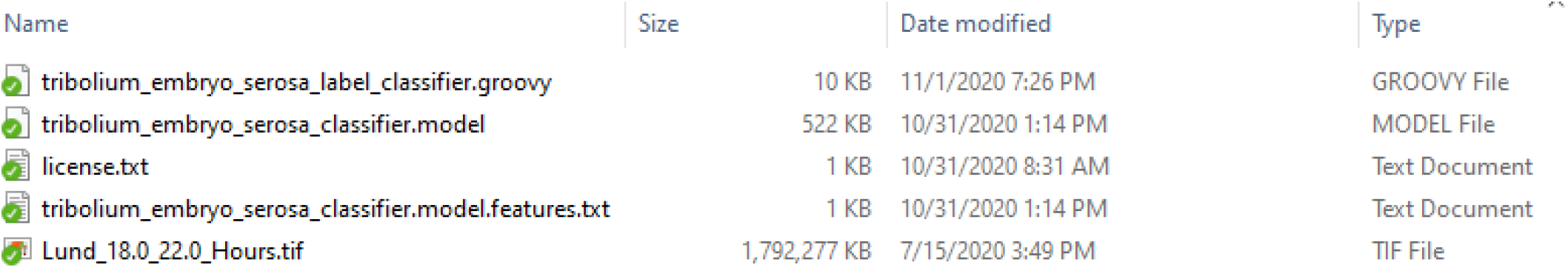
Supplementary material: *Tribolium* data + scripts for the neighborhood relationship classification

**Figure M3:**
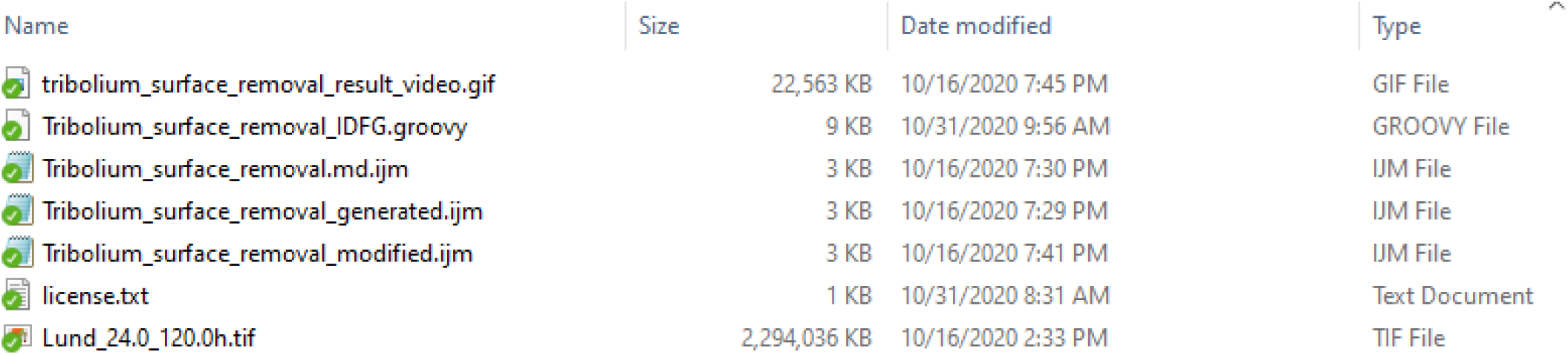
Supplementary material: *Tribolium* data + scripts for the serosa removal example

**Figure M4:**
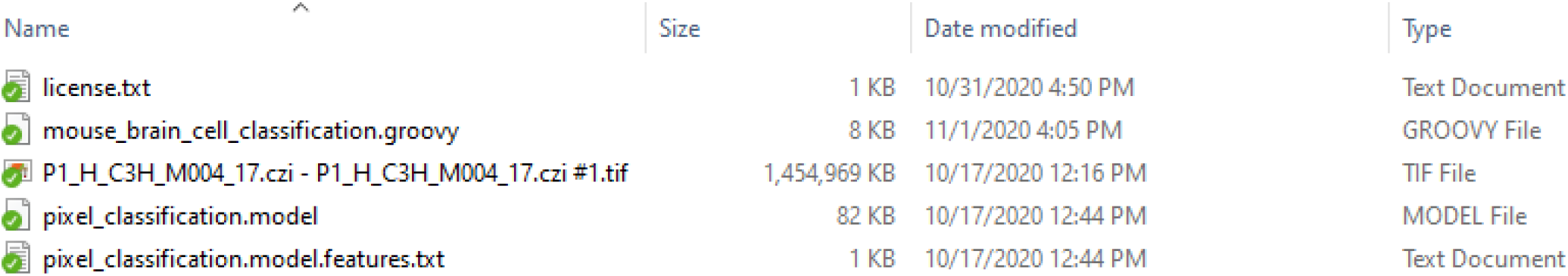
Supplementary material: Mouse brain data + scripts

## Footnotes

1 https://clij.github.io/usage-miner/

2 https://clesperanto.net

## Notes

https://clij.github.io/assistant

https://doi.org/10.5281/zenodo.4276076

### Competing Interest Statement

Research in the laboratories of D.P.P. is funded in part by Takeda Pharmaceuticals International.

